# LRRK2 regulates synaptic function through modulation of actin cytoskeletal dynamics

**DOI:** 10.1101/2022.10.31.514622

**Authors:** Giulia Tombesi, Shiva Kompella, Giulia Favetta, Chuyu Chen, Marta Ornaghi, Yibo Zhao, Ester Morosin, Martina Sevegnani, Adriano Lama, Antonella Marte, Ilaria Battisti, Lucia Iannotta, Nicoletta Plotegher, Laura Civiero, Franco Onofri, Britta J Eickholt, Giovanni Piccoli, Giorgio Arrigoni, Dayne Beccano-Kelly, Claudia Manzoni, Loukia Parisiadou, Elisa Greggio

## Abstract

Parkinson’s disease (PD) is a multisystemic disorder that manifests through motor and non-motor symptoms. Motor dysfunction is the most debilitating and it is caused by the degeneration of dopamine-producing neurons in the substantia nigra pars compacta (SNpc). Increasing evidence suggests that synapse dysfunction precedes neuronal loss by years. Still, early synaptic alterations in PD remain poorly understood.

Here we integrate literature meta-analysis, proteomics and phosphoproteomics with biochemical, imaging and electrophysiological measurements in neurons and brains from knockout and knockin *Lrrk2* mouse models, as well as human iPSC-derived neurons lacking LRRK2. We demonstrate that phosphorylation of LRRK2 at Ser935 and of RAB proteins is induced by brain-derived neurotrophic factor (BDNF) stimulation in differentiated SH-SY5Y cells and primary mouse neurons. Affinity-purification coupled with mass spectrometry (AP-MS/MS) revealed a significant remodelling of the LRRK2 interactome following BDNF treatment, with enhanced association of LRRK2 to a network of actin cytoskeleton-related proteins. Gene-ontology analyses of both literature-curated LRRK2 interactors and phospho-proteome from striatal tissues with elevated LRRK2 activity (G2019S knockin mice) highlight synapse-actin remodelling as major affected pathways.

We further observed that loss of LRRK2 impairs BDNF signaling and alters postsynaptic density architecture. One month-old *Lrrk2* knockout mice display structural alterations in dendritic protrusions, a phenotype that normalizes with age. In human iPSC-derived neurons, BDNF enhances the frequency of miniature excitatory post-synaptic currents (mEPSC) in wild-type cells, an effect that is abolished in the absence of LRRK2.

Taken together, our study discloses a critical role of LRRK2 in BDNF-dependent synaptic modulation and identifies the synaptic actin cytoskeleton as a convergent site of LRRK2-associated pathophysiological processes in PD.

## Introduction

Synaptic damage and connectome dysfunction are emerging as early pathological events preceding neurodegeneration and the manifestation of clinical symptoms in multiple neurodegenerative disorders, including Parkinson’s disease (PD).^1–5^ PD is an age-related motor disorder for which no cure is available. The motor symptoms typically manifest as a consequence of the progressive loss of the dopaminergic neurons of the Substantia Nigra pars compacta (SNpc) projecting to the striatum.^6^ The degeneration of the nigrostriatal projections precedes the loss of the dopaminergic cell bodies in the SNpc and synaptic failure may be the triggering cause of axonal deterioration.^5,7^ Pre- and postsynaptic alterations were described in PD animal models.^5^ For example, PD pathology was reported as associated with an evident striatal spine loss, that appeared to be correlated with the degree of dopamine denervation prevalently due to glutamate excitotoxicity in striatal neurons.^8,9^

Accumulating evidence indicate that more than half of the causative genes and risk factors for PD have a function at the synapse.^10^ Mutations in *LRRK2* gene represent the most common cause of autosomal dominant late onset familial PD and variations around the *LRRK2* locus increase lifetime risk for PD.^11^ Pathogenic mutations are clustered into the enzymatic core of the LRRK2 protein, composed by a Roc/GTPase and a kinase domain, bridged by a COR scaffold.^12–14^ The most frequent mutation (G2019S) located in the kinase domain results in a protein with a gain of kinase activity, associated with increased cellular toxicity.^12^ Previous findings from our team and other laboratories suggested that LRRK2 sits at the crossroads between cytoskeletal dynamics and vesicular trafficking.^15–17^ LRRK2 activity seems to be modulated by interactions with a diverse array of cytoskeletal and vesicle-associated proteins, and it regulates membrane-trafficking events *via* phosphorylation of a subset of Rab GTPases, including RAB3, RAB8, RAB10, RAB12, RAB35 and RAB43 (reviewed in ^15,18,19^). LRRK2 subcellular localization is dynamically controlled by phosphorylation/dephosphorylation of a cluster of N-terminal serine residues (e.g. Ser935 and Ser910), which determines 14-3-3 protein binding.^20^ Phosphorylation of LRRK2 Ser935 and Ser910 is regulated by multiple kinases, including casein kinase 1 alpha (CK1a), IKKs and protein kinase A (PKA),^21–23^ and serves as a scaffold for 14-3-3 protein binding and LRRK2 subcellular localization.^20,24^ In immune cells, extracellular signals such as Toll-like receptor 4 (TLR4) agonists stimulate LRRK2 Ser935 and Ser910 phosphorylation to positively modulate immune-related responses.^21,25^ In contrast, the extracellular signals that orchestrate Ser935 phosphorylation enhancing LRRK2 activity in neurons are still unknown.

The ability of the neurons to relocate LRRK2 within or nearby the synapse could be particularly important as LRRK2 was suggested to provide the scaffold for the assembly of the signalling cascade effectors to sustain synaptic functions.^15,26,27^ In particular, LRRK2 has been shown to regulate synaptic vesicles cycling at the presynaptic site through interaction and phosphorylation of a panel of pre-synaptic proteins,^28^ such as NSF^29^, synapsin I,^30^ EndophylinA,^31^ synaptojanin-1,^32^ and DNAJC6/Auxilin.^33^ Interestingly, recent literature supports a role for LRRK2 in the dendritic spine compartment, where it was suggested to influence AMPA receptor trafficking,^34,35^ spine morphology and functionality in developing brains.^36–38^

Here, starting from the observation that brain derived neurotrophic factor (BDNF) stimulates LRRK2 Ser935 in neurons, we used unbiased protein-protein interaction (PPI) and phosphoproteomic investigations to disclose a novel function of LRRK2 in regulating actin-dependent synaptic dynamics in complementary neuronal models.

## Results

### BDNF treatment rapidly increases LRRK2 phosphorylation at Ser935

Neurotrophic factors plastically shape neuronal activity by binding to their cognate receptor tyrosine kinases.^39^ Brain-derived neurotrophic factor (BDNF) is particularly important to modulate neurotransmission in the cortico-striatal circuitry.^40^ This circuitry is relevant for PD and presents with the highest LRRK2 expression in comparison with the rest of the brain.^41,42^ To address whether LRRK2 activity is stimulated by BDNF, we prepared primary mouse cortical neurons from C57BL/6J wild-type mice and stimulated them with 100 ng/ml of BDNF at days *in vitro* 14 (DIV14) when synapses are fully functional (Fig. 1A) and LRRK2 expression is detectable.^43^ Western blot analysis revealed a rapid and transient increase of Ser935 phosphorylation after BDNF treatment (Fig. 1B and 1C). Of interest, BDNF failed to stimulate Ser935 phosphorylation when neurons were pretreated with the LRRK2 inhibitor MLi-2, indicating that either LRRK2 kinase activity controls the upstream kinases activated by BDNF or that the binding with MLi-2, a type I inhibitor, locks the kinase in a conformation that favors dephosphorylation/hinders phosphorylation during the stimulus. ^44^

**Figure 1.**
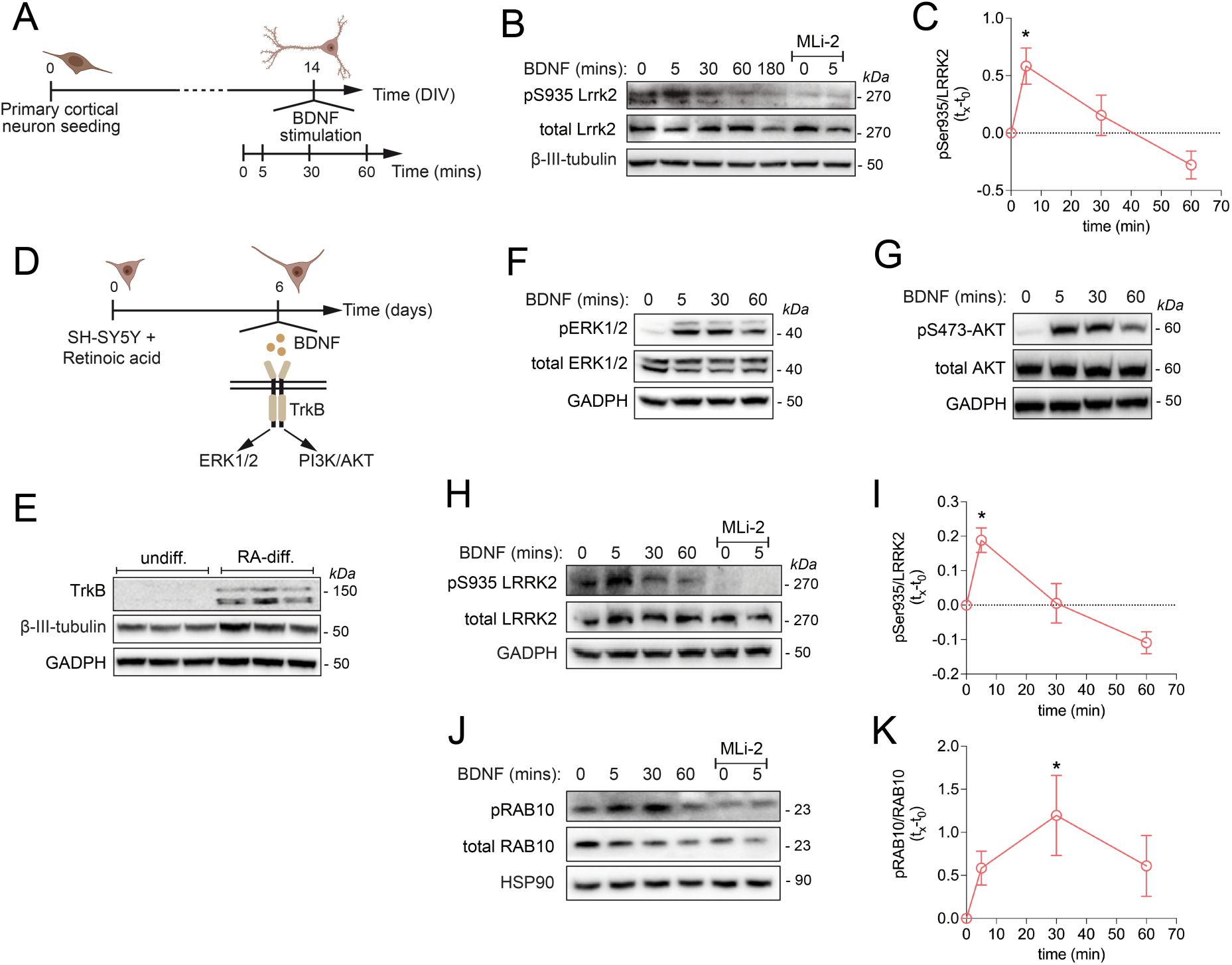
LRRK2 phosphorylation at ser935 rapidly and transiently increases upon BDNF stimulation in primary mouse cortical neurons and differentiated SH-SY5Y cells. **(A)** Schematic representation of the experimental setting of (B). **(B)** Phospho-Ser935 and total Lrrk2 protein levels of primary cortical neurons at DIV14 treated with 100 ng/mL BDNF for 0, 5, 30, 60 180 mins. MLi-2 was used at 500 nM for 90 mins to inhibit Lrrk2 kinase activity. **(C)** Quantification of n=6 independent experiments of (B). One-way ANOVA (\*\*\**P* < 0.001), Dunnett’s multiple comparison test (t=0 compared with different time points, \**P* < 0.05 t=0 vs. t=5’). **(D)** Schematic representation of the experimental setting of (F) and (G). **(E)** TrkB protein levels in undifferentiated or retinoic acid-differentiated SH-SY5Y cells. **(F)** Phospho Thr202/185 ERK1/2, total ERK1/2 and **(G)** phospho Ser473 AKT, total AKT protein levels of retinoic acid-differentiated SH-SY5Y cells stimulated with 100 ng/mL BDNF for 0, 5, 30, 60 mins. **(H)** Phospho-Ser935 and total LRRK2 protein levels of retinoic acid-differentiated SH-SY5Y cells stimulated with 100 ng/mL BDNF for 0, 5, 30, 60 mins. MLi-2 was used at 500 nM for 90 mins to inhibit LRRK2 kinase activity. **(I)** Quantification of (H) (n=4 independent experiments). One-way ANOVA (\*\**P* < 0.01), Dunnett’s multiple comparison test (\**P* < 0.05 t=0 vs. t=5’). **(J)** Phospho-T73 and total RAB10 protein levels of retinoic acid-differentiated SH-SY5Y cells stimulated with 100 ng/mL BDNF for 0, 5, 30, 60 mins. MLi-2 was used at 500 nM for 90 mins to inhibit LRRK2 kinase activity. **(K)** Quantification of (**J**) (n=4 independent experiments). One-way ANOVA (*P* > 0.5), Dunnett’s multiple comparison test (\**P* < 0.05 t=0 vs. t=30’).

To gain mechanistic insights into the effects of BDNF stimulation on LRRK2 function, we differentiated neuroblastoma SH-SY5Y cells with 10 µM retinoic acid (RA) treatment for 6 days^45^ and measured BDNF response by monitoring Tropomyosin receptor kinase B (TrkB) levels and phosphorylation of AKT and ERK1/2, two major downstream effector pathways of BDNF/TrkB signaling (Fig. 1D).^46^ Expression of TrkB was undetectable in undifferentiated SH-SY5Y cells but was induced upon RA differentiation (Fig. 1E). Accordingly, differentiated SH-SY5Y cells respond to 100 ng/ml BDNF stimulation, as evidenced by increased phosphorylation of ERK1/2 and AKT (Fig. 1F-G). LRRK2 expression, low in undifferentiated cells, also increased upon differentiation (Fig. S1A), indicating that this is a suitable neuronal model to study endogenous LRRK2. Notably, BDNF stimulation enhanced Ser935 phosphorylation with kinetics comparable with that observed in primary neurons (Fig. 1H and 1I). To determine whether BDNF-induced Ser935 phosphorylation reflects LRRK2 kinase activation, we assessed phosphorylation of RAB10 at Thr73, an established LRRK2 substrate. As illustrated in Fig. 1J and 1K, BDNF treatment significantly increased RAB10 phosphorylation, supporting the role of BDNF as a physiological activator of LRRK2.

### BDNF stimulates LRRK2 interaction with drebrin, an actin cytoskeletal-associated protein enriched at the postsynapse

To clarify the implications of BDNF-mediated LRRK2-activation on neuronal function, we used affinity purification coupled with mass spectrometry (AP-MS/MS). As the yields of endogenous LRRK2 purification were insufficient for AP-MS/MS analysis, we generated polyclonal SH-SY5Y cells stably expressing GFP-LRRK2 wild-type or GFP control. Notably, in this model, Ser935 phosphorylation showed only a modest increase upon BDNF treatment (Fig. S1B), and prolonged stimulation led to a reduction of Ser935 phosphorylation below baseline levels (Fig. 1B–C and Fig. 1H-I). We hypothesized that this apparent limited response could be due to high basal levels of Ser935 phosphorylation, approaching a ceiling effect under resting conditions. This is consistent with previous mass spectrometry analyses reporting that Ser935 is nearly fully phosphorylated (∼100%) in the brain under basal conditions.^47^ To test whether high basal phosphorylation masks BDNF-induced increases, we pretreated the cells with MLi-2 for 90 minutes to lower Ser935 phosphorylation. After inhibitor washout, cells were stimulated with BDNF at multiple time points. In these conditions, BDNF robustly increased both Ser935 and RAB phosphorylation, peaking at 5–15 minutes and returning to baseline by 60–180 minutes (Fig. S1C). Notably, while the magnitude of pSer935 phosphorylation at the peak was similar to untreated controls - supporting the idea that Ser935 is near saturation under basal conditions - RAB phosphorylation levels significantly exceeded those of untreated cells. This indicates that, unlike Ser935, RAB phosphorylation remains far from saturated under basal conditions and serves as a more dynamic readout of LRRK2 kinase activity. These data further support BDNF as a physiological activator of LRRK2 and validate GFP-LRRK2 SH-SY5Y cells for interrogating stimulus-induced changes in LRRK2 signaling and protein interactions.

Next, we used GFP-trap purification to isolate GFP or GFP-LRRK2 from RA-differentiated cells unstimulated or stimulated with BDNF for 15 mins prior to MS analysis. The unstimulated LRRK2 interactome (GFP-LRRK2 vs. GFP) is large (207 significant hits) and consistent with previous computational analysis of LRRK2 PPIs (Fig. 2A).^48,49^ In particular, we confirmed interactions with known LRRK2 binders including 14-3-3 proteins, cytoskeletal proteins and protein translation factors (Supplementary.xls table 1). Since BDNF is synaptically released and influences pre- and post-synaptic mechanisms, we used SynGO^50^ to search for LRRK2 interactors that are enriched at the synapse. About one-third (35%) of LRRK2 PPI under unstimulated conditions are SynGO annotated genes; among them, 33% are presynaptic proteins and 66% are post-synaptic proteins (Fig. 2B). The most significant enriched categories were pre- and post-synaptic ribosome and post-synaptic actin cytoskeleton (Fig. 2B).

**Figure 2.**
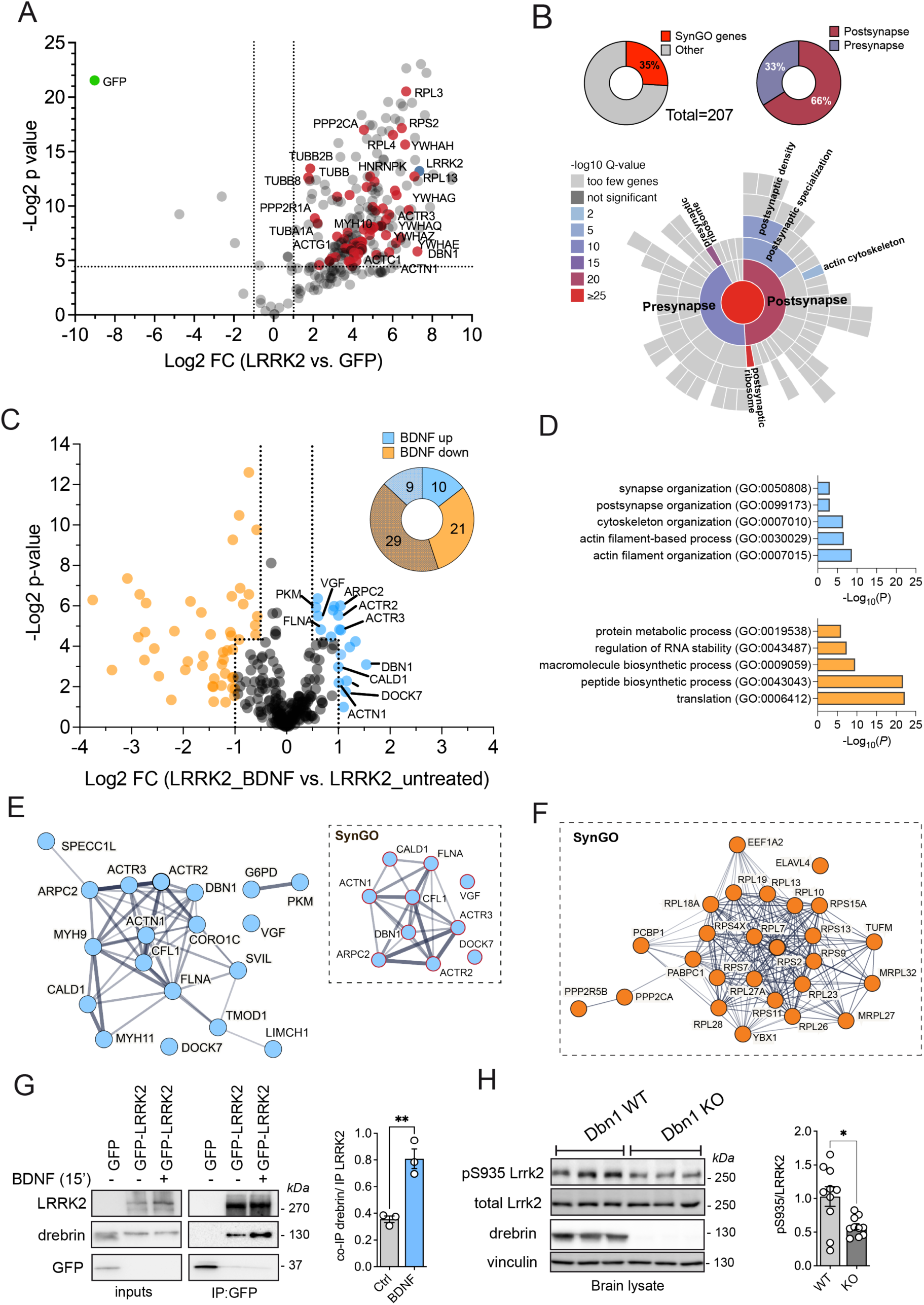
BDNF promotes LRRK2 interaction with post-synaptic actin cytoskeleton components. **(A)** Volcano plot of GFP-LRRK2 versus GFP enriched interactors from differentiated SH-SY5Y cells (n=6 replicates: n=3 independent experiments each with 2 technical replicates). Interactors that were considered for SynGO analysis have adjusted *P* value < 0.05 and FC > 2. (red dots). **(B)** Donut charts with % of SynGO genes among the total 207 LRRK2 interactors (A) and with % of presynaptic and postsynaptic proteins among SynGO genes. SynGO-CC terms visualized with a sunburst plot showing significant categories (below). **(C)** Volcano plot of GFP-LRRK2 +/-BDNF enriched interactors from differentiated SH-SY5Y cells (n=6 replicates: n=3 independent experiments each with 2 technical replicates). Interactors selected for pathway enrichment analysis fall into two categories: (i) adjusted *P* value < 0.05 and FC > 0.5 or FC < −0.5, or (ii) |FC| > 1 regardless of the *P* value. Proteins with increased interaction upon BDNF stimulation are blue-labeled, proteins with decreased interaction are orange-labeled. Donut chart showing the number of SynGO annotated genes versus non synaptic proteins in BDNF-up versus BDNF-down interactors (top right). **(D)** G:profiler g:GOSt pathway enrichment analysis showing the top 5 enriched biological processes (BP) categories of BDNF-increased interactors (top, blue bar graph) versus BDNF-decreased interactors (bottom, orange bar graph). GO analysis was performed with g:Profiler on 04/14/2025 (version e112_eg59_p19_25aa4782, database updated on 03/02/2025); enriched terms determined using g:SCS correction and size terms set to <500 to increase specificity. **(E)** Protein-Protein Interaction Networks built with STRING (https://string-db.org/) of all BDNF(+) interactors and SynGO BDNF-increased interactors (inset) and **(F)** of SynGO BDNF-decreased interactors. **(G)** Validation of increased drebrin:LRRK2 interaction upon BDNF treatment (15 mins) and quantification (one sample t-test, *P*<0.01). Samples are the same analyzed in the three rounds of MS (A). **(H)** Western blot analysis of brain samples from WT and *Dbn1* (drebrin) KO mice (2 month old) assessing Lrrk2 Ser935. Differences between the two genotypes have been evaluated using Student’s t-test (significance \**P*<0.05 pSer935), *n*=10 animals per genotype.

**Table 1.**
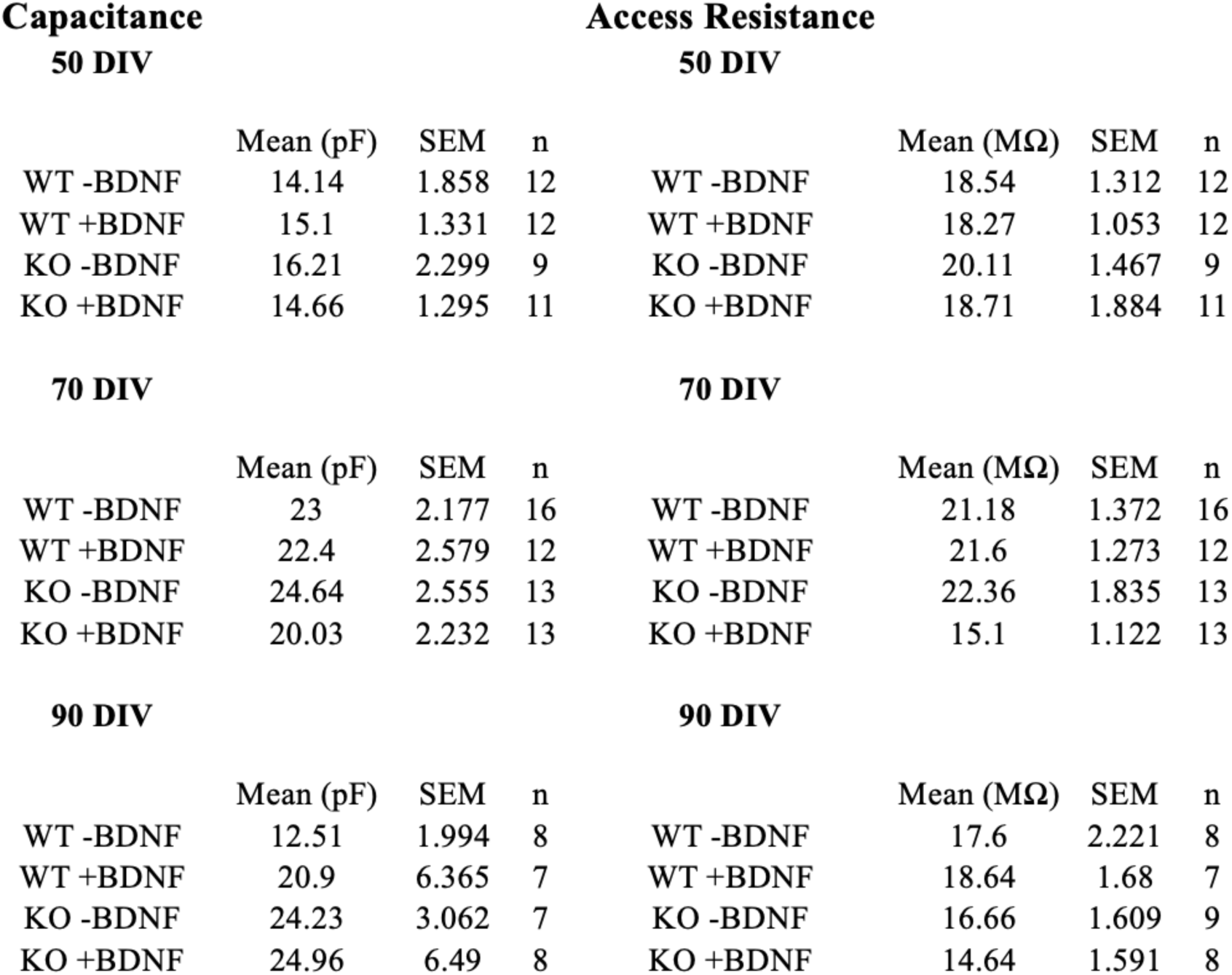
Capacitance and access resistance of LRRK2 KO and WT cortical neurons at 50, 70 and 90 DIV with or without BDNF stimulation.

We next compared unstimulated vs. BDNF-stimulated LRRK2 interactome. As shown in Fig. 2C, 15 minutes of BDNF stimulation reshapes LRRK2 PPI, with a group of interactors increasing binding with LRRK2 (blue dots) and another group decreasing binding (orange dots). Inclusion criteria were: (i) *P* value < 0.05 and fold change (FC) > 0.5 or FC < −0.5, or (ii) absolute fold change (|FC|) > 1 regardless of the *P* value. In this way, we increased the discovery power for gene ontology analysis. SynGO gene ontology revealed that 10 out of 19 hits with increased binding after BDNF [BDNF(+)] and 21 out of 50 hits with decreased binding [BDNF(-)] are synaptic proteins (Fig. 2C inset). Among synaptic interactors, the BDNF(+) binders are enriched in actin-cytoskeleton-related GO:BP categories while the BDNF(-) binders are enriched in protein translation GO:BP categories (Fig. 2D). Furthermore, 80% of synaptic BDNF(+) LRRK2 binders (DBN1, ARPC2, ACTR2, ACTR3, CFL1, CALD1, ACTN1, FLNA) and 62% of synaptic BDNF(-) binders (RPS2, RPS4X, RPS7, RPS9, RPS11, RPS13, RPS15A, RPL13, RPL23, RPL26, RPL27A, TUFM, EEF1A2, PCBP1) form functional and physical interaction networks (Fig. 2E-F).

To validate these findings, we performed western blot analysis on the AP samples analyzed by MS. We selected drebrin, encoded by *DBN1*, because it sits at the center of the BDNF(+) network (Fig. 2E, inset SynGO) and, in DIV14 primary neurons, it colocalizes with PSD95, a postsynaptic marker (Fig. S2A), confirming previous observations.^51^ As predicted, BDNF treatment increases LRRK2:drebrin interaction by approximately twofold (Fig. 2G and Fig. S2B). To further characterize this interaction, we performed co-immunoprecipitation in HEK293T cells co-expressing Flag-LRRK2 and either full-length YFP-drebrin, its N-terminal (1-256 aa), or C-terminal (256-649 aa) fragments. These experiments confirmed a direct interaction between LRRK2 and drebrin, with the N-terminal domain appearing to mediate the majority of the binding (Fig. S2C). Notably, this region contains both the ADF-H (actin-depolymerizing factor homology) domain and a coiled-coil motif known to associate with actin.^51,52^ Finally, we observed a significant decrease in Ser935 phosphorylation in *Dbn1* KO mouse brains (Fig. 2H). This finding further strengthen the notion that drebrin forms a complex with LRRK2 that is important for its activity, e.g. upon BDNF stimulation.

We conclude that BDNF stimulation engages LRRK2 into actin cytoskeleton dynamics via its interaction with actin-regulatory proteins. While SH-SY5Y cells may only contain sparse synapses,^53^ the enrichment of this BDNF-dependent protein network at synapses indicates that this site might be relevant for LRRK2 function. Our findings also suggest that a pool of LRRK2 binds protein translation factors under unstimulated conditions, which redistributes toward actin-rich domains upon BDNF-TrkB signaling.

### Convergence on actin and synaptic processes in literature-curated LRRK2 interactome and LRRK2 G2019S striatal phospho-proteome

We recently investigated the molecular environment surrounding LRRK2 specifically by constructing the protein-protein interaction (PPI) interactome around LRRK2 based on PPIs derived from peer reviewed literature. After filtering the LRRK2 PPIs for reproducibility (keeping only those interactions that have been reported at least twice in literature or at least replicated with 2 different interaction detection methods) we obtained the interactions linking across the LRRK2 interactors thus effectively building the LRRK2 protein interaction network (LRRK2_net_) which describes the protein milieu around LRRK2 in the cellular context (see Zhao *et al.*^54^ for details). The LRRK2_net_ was topologically analyzed to identify the portions (clusters) of the network that are more densely connected locally in comparison with the overall connectivity of the entire network as these topological clusters are likely to represent functional units.^55^ Among the 11 topological clusters identified in Zhao *et al*.^54^, we further elaborated on one cluster in particular, containing 41 genes (Fig. 3A) and found that it exhibited significantly GO-BP enriched terms into semantic categories related to: actin polymerization (16 GO-BP terms), synaptic vesicles (5 GO-BP terms), postsynaptic actin-cytoskeleton (2 GO-BP terms), lamellipodium (6 GO-BP terms), cell-neuron projection/size (2 GO-BP terms), DNA recombination/repair (4 GO-BP terms) and uncategorized (6 GO-BP terms) (Fig. 3B and Supplementary.xls table 2). Out of the 41 proteins forming the cluster, 23 are proteins associated with the actin cytoskeleton including the 4 larger nodes (where the dimension of the nodes is directly proportional to the node degree/connections within the network): DBN1/drebrin, CAPZA2 and LIMA whose association with LRRK2 was identified in Meixner *et al.*^56^ and IQGAP1 identified in Liu *et al.*^57^ (Fig. 3C). Given the overrepresentation of synapse-related GO-BP terms, we further performed GO functional enrichment using SynGO and mapped 16 synaptic proteins within the cluster (Fig. 3C). Remarkably, DBN1/drebrin is a key protein in the cluster, as it engages in multiple interactions (16 edges) and sits at the interface between the nodes involved in GO:BPs functions related to the actin cytoskeleton (blue-colored nodes) and to the synapse (red-colored nodes) (Fig. 3C and Fig. S3).

**Figure 3.**
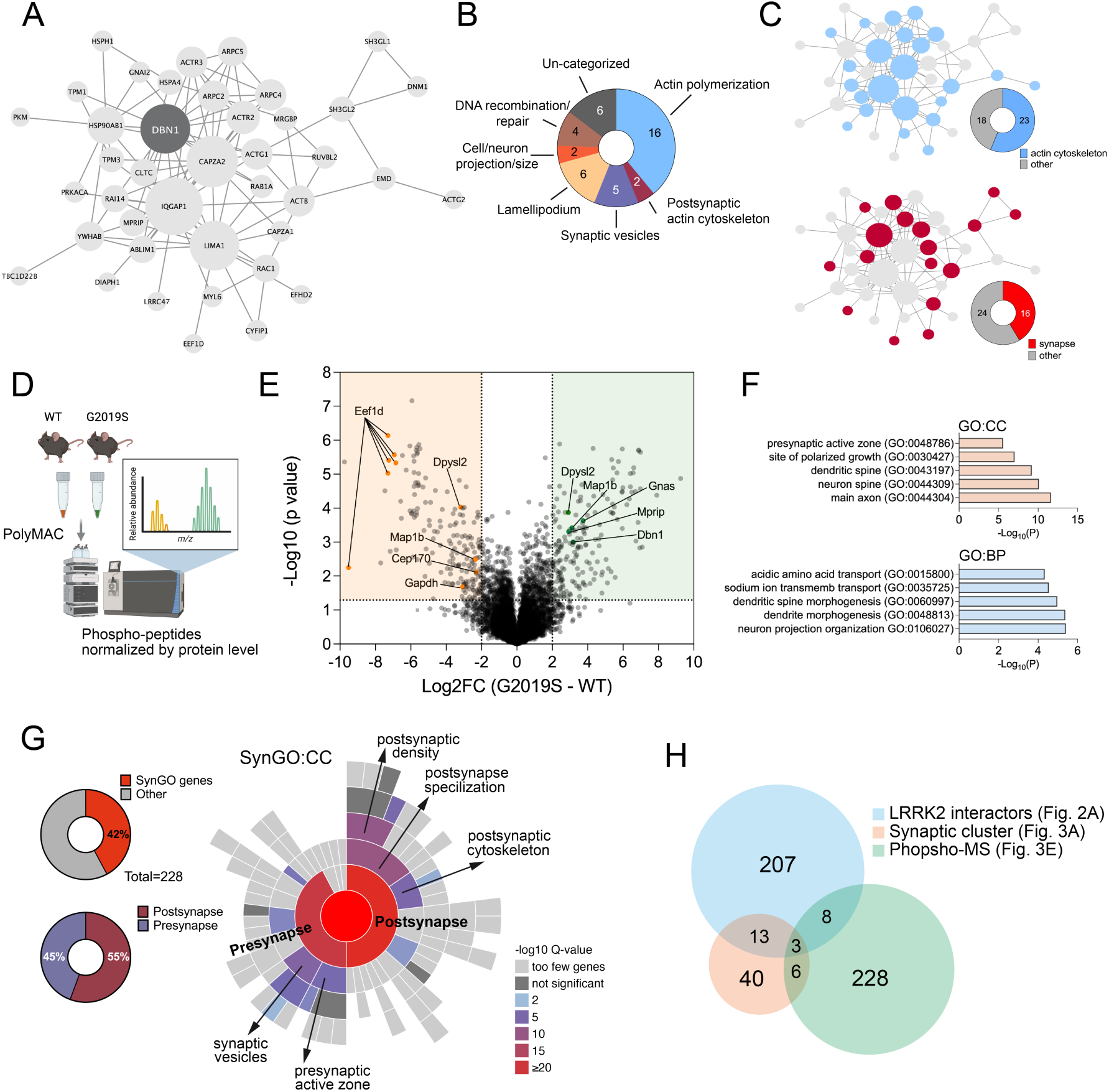
Convergence on actin and synaptic processes in literature-curated LRRK2 interactors and the LRRK2 G2019S striatal phospho-proteome. **(A)** Network graph showing one of the topological clusters extracted from the LRRK2 protein-protein interaction network. Proteins are represented as nodes while protein-protein interactions are represented as edges. Node size is proportional to the degree centrality: the larger the node, the higher the degree, the more interactions the node has in the network. **(B)** Gene Ontology biological-processes (GO-BPs) enrichment; the analysis was performed using g:Profiler, GO-BPs with term size >500 were discarded as general terms and the remaining terms was clustered based on semantics using the keywords: Postsynaptic (Actin) Cytoskeleton, Lamellipodium, Synaptic Vesicle, Actin Polymerization/Nucleation, DNA Recombination/Repair, Cell/Neuron Projection. **(C)** Upper network graph as in (A) highlighting actin cytoskeletal proteins in blue and the proportion over the total (pie chart) and lower network graph as in (A) highlighting synaptic proteins in red and the proportion over the total (pie chart). (**D**) Experimental design of phospho-proteomic analysis comparing WT and G2019S knockin mouse striata. Phosphopeptide abundances were normalized to total proteins levels. **(E)** Volcano plot of differentially enriched phosphopeptides, expressed as Log2 fold change (FC) between G2019S-KI versus WT. Highlighted peptides correspond to LRRK2 interactors discovered by MS in Figure 2. **(F)** GO enrichment analysis performed with g:Profiler (accessed on 01/23/2025; version e94_eg41_p11), filtering for GO terms with size <200 to avoid overly general categories. The top 5 most significant terms from the Cellular Component (CC) and Biological Process (BP) domains are shown. **(G)** Sunburst plot showing enriched SynGO cellular component (CC) categories and the percentage of SynGO genes among the differentially phosphorylated proteins. **(H)** Venn diagrams showing common and unique hits across the three datasets: GFP-LRRK2 interactomics, synaptic cluster and phospho-MS.

Given the multiple lines of evidence pointing to the synapse-actin couple as a key area for LRRK2 function, and based on the increased LRRK2 activity upon neurotrophin stimulation, we next asked whether LRRK2 kinase activity is also linked to synaptic function. To this aim, we performed an unbiased phospho-proteomic characterization of the striatum, using knockin mice expressing the hyperactive G2019S mutation to probe the contribution of kinase activity. We isolated total striatal proteins from WT and G2019S striata, enriched the phosphopeptides (Fig. 3D) (also see^58^), and normalized the log_2_ intensity of each phosphopeptide to the log_2_ intensity of its corresponding total protein. Comparing WT and G2019S samples, we identified 228 differentially regulated phosphopeptides (*P* value < 0.05 and |Log_2_ fold change (FC)| > 2) (Fig. 3E and supplementary.xls table 3). Gene ontology analysis of differentially phosphorylated proteins, using stringent term size (<200 genes), revealed enrichment of post-synaptic spines and presynaptic active zones (Fig. 3F). A SynGO analysis confirmed enrichment of both pre- and postsynaptic categories, with particular high significance for terms related to the postsynaptic cytoskeleton (Fig. 3G). Despite starting from whole striatal tissue - which contains a mix of different cellular populations - the strong synaptic enrichment of phosphorylation changes further supports the critical role of this compartment in LRRK2 brain function. By merging the three datasets (the LRRK2 interactome in SH-SY5Y cells, the literature-based synaptic/actin cluster and the phosphoproteomic dataset from G2019S striatum), we found three common hits: Dbn1/drebrin, Eef1d and Mprip (Fig. 3H). In particular, phosphorylation of drebrin at Ser339 was 3.7 fold higher in G2019S mice. This site is located in the N-terminal region of drebrin, which exhibited higher affinity for LRRK2 (Fig. 3E and Fig. S2C). To infer the potential physiological relevance of this phosphorylation, we examined its detection frequency in phosphoproteomic studies (67 observations, https://www.phosphosite.org/proteinAction.action?id=2675&showAllSites=true) and assessed its predicted pathogenicity using AlphaMissense.^59^ Although the AlphaFold structural model has low confidence in this region of drebrin, amino acid substitutions at Ser339 are predicted to be non-pathogenic (Fig. S4), supporting the notion that LRRK2-mediated phosphorylation at this site acts as a regulatory modification for drebrin activity.

### BDNF-mediated signalling is impaired in LRRK2 knockout neurons

Having collected multiple lines of evidence linking LRRK2 to actin cytoskeletal proteins enriched at the synapse, and given that BDNF - a neurotrophin known to promote synaptic plasticity - activates LRRK2 (Figure 1), we next asked whether loss of LRRK2 could affect BDNF signalling. To this end, we generated LRRK2 knockout (KO) SH-SY5Y cells with CRISPR/Cas9 editing (Fig. S5 and supplemental file 1), differentiated them with RA and assessed their response to BDNF stimulation. While application of BDNF increased AKT and ERK1/2 phosphorylation in naïve SH-SY5Y, as expected, LRRK2 KO cells exhibited a weaker response (Fig. 4A-B). These results indicate that LRRK2 acts downstream of TrkB and upstream of AKT and ERK1/2, in line with its rapid and transient phosphorylation kinetic upon BDNF stimulation (Fig. 1B-C).

**Figure 4.**
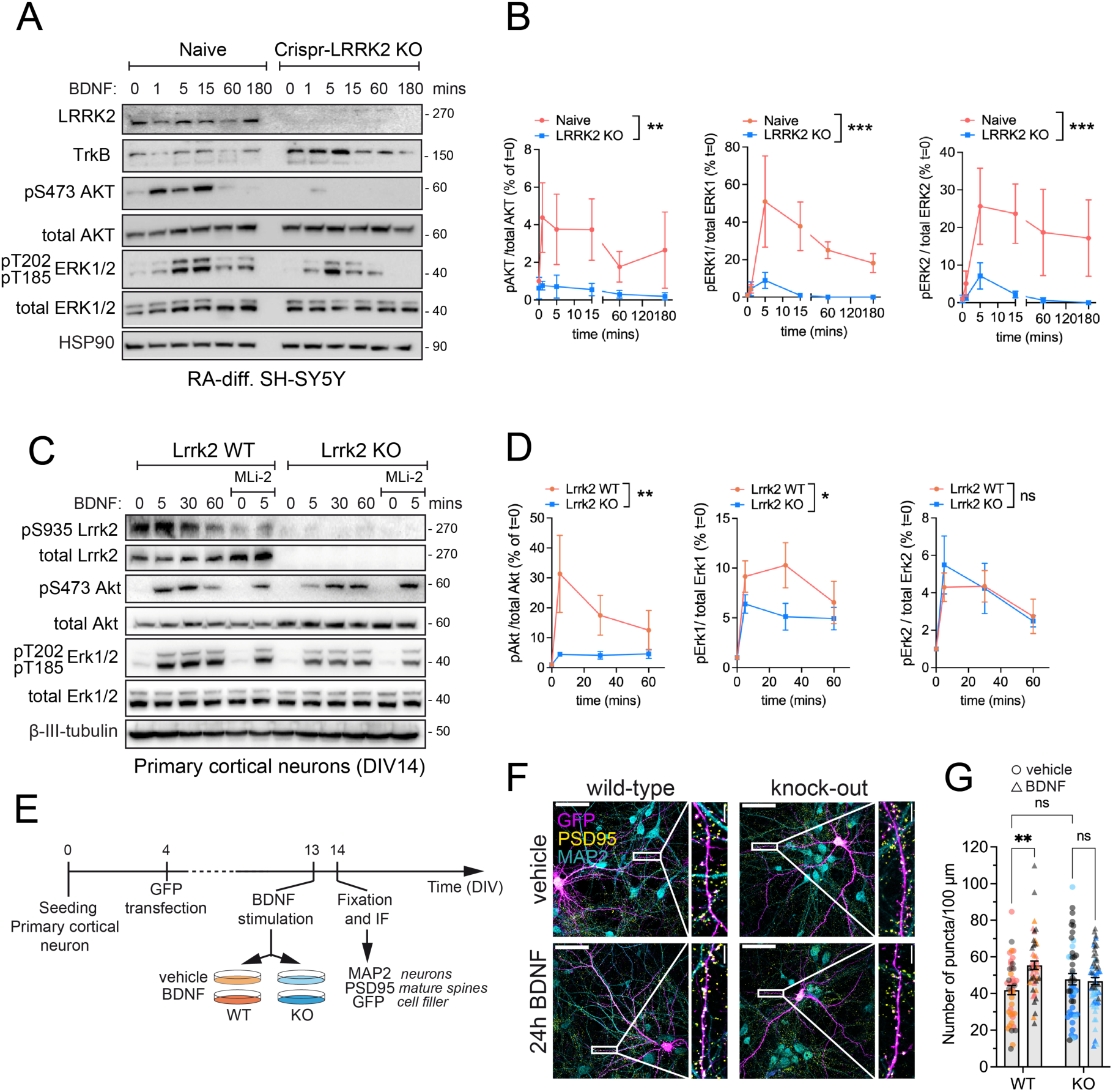
BDNF signaling is impaired in *Lrrk2* knockout neurons. **(A)** Six-day RA**-**differentiated SH-SY5Y naïve and CRISPR-LRRK2 KO cells stimulated with BDNF at different time points (0, 1, 5, 15, 60, 180 mins). Anti-TrkB antibodies were used to confirmed expression of BDNF receptor. To compare BDNF-induced signaling in naïve vs LRRK2-KO cells, phosphorylated Akt (S473) and Erk1/2 (T185/T202) were evaluated. **(B)** Quantification of phosphorylated proteins show from *n*=3 independent differentiation experiments. Two-way ANOVA; phospho-Akt: interaction *P*=0.7079, F (5, 24) = 0.5896; genotype: \*\**P*=0.0014, F (1, 24) = 13.07; time: *P*=0.5610, F (5, 24) = 0.7994. Phospho-Erk1: interaction *P*=0.0783, F (5, 24) = 2.284; genotype: \*\*\**P*=0.0003, F (1, 24) = 17.58; time: \**P*=0.0244, F (5, 24) = 3.174. Phospho-Erk2: interaction: *P*=0.3725, F (5, 24) = 1.128; genotype: \*\*\**P*=0.0008 F (1, 24) = 14.59; time: *P*=0.1356, F (5, 24) = 1.879. **(C)** DIV14 primary cortical neurons from WT vs KO mice stimulated with BDNF at different time points (0, 5, 15, 30, 60 mins) in the presence or absence of LRRK2 inhibitor MLi-2 (90 min, 500 nM). To compare BDNF-induced signaling in WT vs KO neurons, phosphorylated Akt (S473) and Erk1/2 (T185/T202) were evaluated. Phosphorylated LRRK2 was assessed with pS935 antibodies. **(D)** Quantification of phosphorylated proteins show from *n*=9 independent cultures. Two-way ANOVA; phospho-Akt: interaction *P*=0.1186, F (3, 63) = 2.031; genotype: \*\**P*=0.0037, F (1, 63) = 9.101; time: *P*=0.0342, F (3, 63) = 3.069. Phospho-Erk1: interaction *P*=0.3256, F (3, 64) = 1.177; genotype: \**P*=0.0201, F (1, 64) = 5.680; time: \*\*\*\**P*<0.0001, F (3, 64) = 10.04. Phospho-Erk2: interaction *P*=0.8524, F (3, 62) = 0.2622; genotype: *P*=0.7528, F (1, 62) = 0.1001; time: \*\*\**P*=0.0003, F (3, 62) = 7.428. **(E)** Overview of the experimental workflow to induce spine formation/maturation. **(F)** Representative confocal images of primary cortical neurons stimulated with 100 ng/ml of BDNF or vehicle control for 24 hours. GFP has been transfected at DIV4 to fill the neuroplasm and visualize individual dendrites, MAP2 is a neuronal marker and PSD95 is a marker of mature spines. **(G)** Quantification of the number of PSD-positive puncta per unit of length (100 µm). Dots represents individual segments (*n*=20 neurites from *n*=5-6 neurons per replicate) and colors define neuronal cultures prepared in different days from pulled pups (*n*=8 pups per culture per genotype, *n*=3 independent cultures). Two-way ANOVA; interaction \*\**P*=0.0038, F (1, 191) = 8.584; genotype: *P*=0.5833, F (1, 191) = 0.3020; treatment: \**P*=0.0126, F (1, 191) = 6.346. Šídák’s multiple comparisons test: vehicle vs. BDNF (WT) \*\*\**P*=0.0005; vehicle vs. BDNF (KO) *P*=0.9440; Vehicle (WT) vs. vehicle (KO) *P*= 0.3756.

Likewise, *Lrrk2* WT and KO primary cortical neurons at DIV14 were stimulated with BDNF at different timepoints. Similar to differentiated SH-SY5Y cells, *Lrrk2* WT neurons responded to BDNF by rapidly increasing Akt and Erk1/2 phosphorylation, whilst KO neurons exhibited a significantly reduced phosphorylation of Akt and Erk1 (Fig. 4C-D).

Considering that (i) BDNF induces dendritic spine formation, maturation and structural plasticity,^60,61^ (ii) BDNF(+) LRRK2 interactors are actin-related proteins enriched at the postsynapse (Fig. 2) and (iii) loss of LRRK2 causes reduced phosphorylation of BDNF/TrkB downstream kinases AKT and ERK1/2 (Fig. 4A-D), we evaluated BDNF-induced PSD95 sites in the presence or absence of LRRK2, as a potential process affected by LRRK2 deletion. Primary cortical neurons were transfected with GFP filler at DIV4 and treated with BDNF for 24 h at DIV13 following previous protocols^62,63^ (Fig. 4E). As shown in figure 4F-G, BDNF treatment significantly increased the density of PSD95-positive puncta in WT cultures (*P* < 0.001). In contrast, *Lrrk2* KO neurons did not exhibit increased PSD95 puncta upon BDNF. These findings further support a role for LRRK2 within the BDNF signalling cascade and show that the absence of LRRK2 impairs the remodelling of dendrite protrusions in response to BDNF stimulation.

### Morphological Alterations of Dendritic Protrusions in Young Lrrk2 Knockout Mice

Based on *in vitro* results indicating that BDNF stimulates LRRK2 activity and promotes its recruitment to the actin-cytoskeleton, enhancing the number of PSD95-positive dendritic protrusions, we next investigated whether loss of *Lrrk2* results in dendritic spine defects in the mouse brain. During development, dendritic spines are highly dynamic, with a rate of extension and retraction that allows proper and accurate synapse formation and neuronal circuit assembly.^64^ By the adulthood, dendritic spines formation, motility, and pruning decrease reaching a relatively stable number of protrusions.^65^ Since spines dynamics vary according to the stage of brain development, we performed a longitudinal analysis of young (1 month-old), adult (4 month-old) and aged (18 month-old) *Lrrk2* WT and KO mice in order to capture any age-dependent defect (Fig. 5A). We focused on the dorsal striatum, a region highly enriched in SPNs that receives excitatory afferents from the cortex and dopaminergic modulation from the SNpc. This region is relevant in the prodromal stages of PD neurodegeneration^66^ and expresses high levels of LRRK2.^41,67^ Moreover, several studies showed a key role of LRRK2 in shaping the structure and function of excitatory synapses in the striatum.^36,37,68,69^ We evaluated the number and morphology of dendritic spines using Golgi-Cox staining (Fig. 5B). The total number of protrusions varied across ages, but there were no differences between the two genotypes (Fig. 5C). Instead, the width and the length of the protrusions were reduced in 1 month-old Lrrk2 KO striata with respect to WT. Specifically, the average neck height was 15% shorter and the average head width was 27% smaller, indicating that the protrusions are smaller in *Lrrk2* KO brains. No differences were observed at 4 and 18 months of age (Fig. 5C). We then classified the morphology of the protrusions into four categories: filopodia and thin protrusions (immature spines) and mushroom and branched protrusions (mature spines). One month-old *Lrrk2* KO animals exhibit a reduced number of filopodia and an increased quantity of thin protrusions, suggesting that loss of Lrrk2 may favor this transition. Instead, the proportion of mature spines (mushroom and branched) remained unaltered. No differences were observed in older mice (Fig. S6).

**Figure 5.**
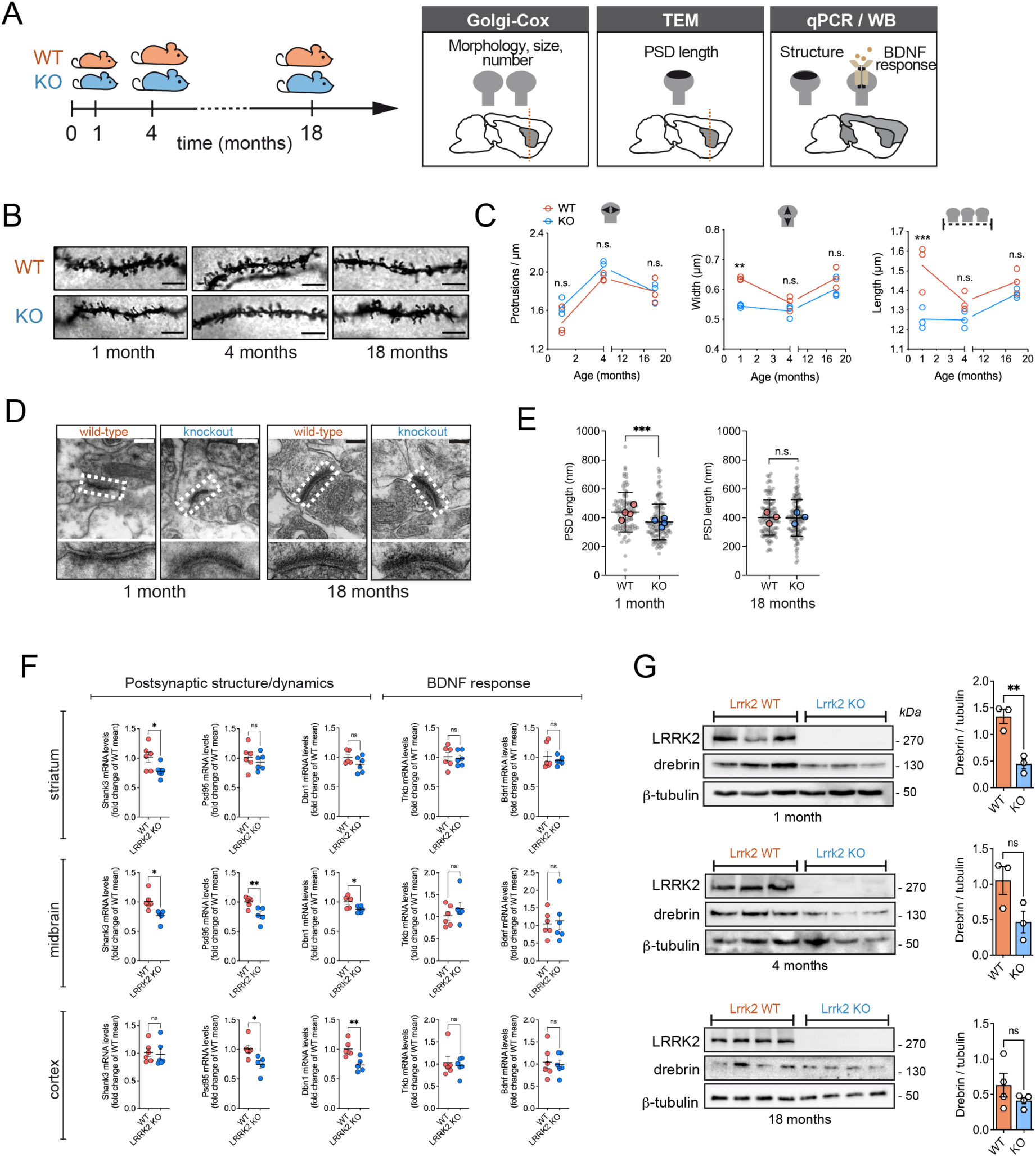
Postsynaptic structural changes in young *Lrrk2* knockout mice. **(A)** Overview of experimental design. **(B)** Representative images of neurite segments from Golgi-Cox stained neurons of dorsal striatum. Scale bar: 3 µm. **(C)** Quantification of average spine number (top), width (middle) and length (bottom) of *n*=3 animals per group (same segments analyzed in B). Statistical significance was determined by two-way ANOVA with Šídák’s multiple comparisons test. Number of protrusions: interaction *P*=0.3820, F (2, 12) = 1.044; age: \*\*\*\**P*<0.0001, F (2, 12) = 27.12; genotype: *P*=0.0840, F (1, 12) = 3.550; WT vs. KO (1 month) *P*=0.2013; WT vs. KO (4 months) *P*=0.5080; WT vs. KO (18 months) *P*>0.9999. Width: interaction *P*=0.0815, F (2, 12) = 3.112; age: \*\*\**P*=0.0004, F (2, 12) = 16.14; genotype: \*\*\**P*=0.0007, F (1, 12) = 20.77; WT vs. KO (1 month) \*\**P*=0.0017; WT vs. KO (4 months) *P*=0.4602; WT vs. KO (18 months) *P*=0.2489. Length: interaction \**P*=0.0345, F (2, 12) = 4.514; age: \**P*=0.0203, F (2, 12) = 5.488; genotype: \*\*\**P*=0.0006, F (1, 12) = 21.53; WT vs. KO (1 month) \*\*\**P*=0.0008; WT vs. KO (4 months) *P*=0.2875; WT vs. KO (18 months) *P*=0.5948. **(D)** Representative transmission electron microscopy (TEM) micrographs of striatal synapses from 1-month and 18-month old WT vs KO mouse brain slices. Scale bar: 200 nm. **(E)** Quantification of post-synaptic density (PSD) length. Graphs show the length of individual synapses (grey dots) and the average PSD length per animal (colored dots). Statistical significance has been calculated with Student’s t-test: 1 month-old mice (*n*=4 mice, 95 synapses WT, 118 synapses KO, \*\*\**P*=0.0003); 18-month old mice (*n*=3 mice, 108 synapses WT, 119 synapses KO, *P*=0.8502). **(F)** Quantitative PCR of *Bdnf, TrkB, Psd95, Shank3* and *Dbn1* mRNA expression in striatum, cortex and midbrain form *n*=6 *Lrrk2* WT and *n*=6 *Lrrk2* KO one-month old mice. Statistical significance has been calculated with Student’s t-test. (**G**) Western blot analysis of whole brain samples from the same Lrrk2 WT and KO mice where Golgi-Cox staining has been performed. The reduction in drebrin content is significant in Lrrk2 KO mice at 1 month of age. Differences between the two genotypes have been evaluated using Student’s t-test (significance \*\**P*<0.01), *n*=3-4 animals per genotype per age.

Since the overall width/height of *Lrrk2* KO protrusions is reduced (Fig. 5C), we next analyzed the postsynaptic ultrastructure using electron microscopy. To increase the experimental power, we used a separate group of animals. We measured the postsynaptic density (PSD) length as it directly correlates with the amount of postsynaptic receptors and signaling proteins, thus providing an indication of the synaptic strength (Fig. 5D). PSD is significantly shorter in 1 month-old *Lrrk2* KO animals compared to WT (*P* < 0.001), while no differences in PSD length are observed in 18 month-old KO mice (Fig. 5E). Taken all these data together, we conclude that subtle defects in the postsynaptic density are present in the striatum of postnatal *Lrrk2* KO mice, but these abnormalities appear to normalize with age.

To assess whether the loss of *Lrrk2* in young mice leads to reduced dendritic protrusion size and postsynaptic density through alterations in BDNF-TrkB signaling or postsynaptic scaffold expression, we measured transcript levels of *Bdnf*, *TrkB*, *Dbn1, Psd95* and *Shank3*. Neither *Bdnf* nor *TrkB* expression is altered in the striatum, cortex or midbrain of 1-month old *Lrrk2* KO as compared to WT mice (Fig. 5F), supporting a mechanism whereby LRRK2 acts downstream of BDNF signaling. Instead *Dbn1*, *Psd95* and *Shank3* expression levels are reduced in *Lrrk2* KO mice across most brain regions analyzed, with the most pronounced decrease observed in the midbrain (Fig. 5F).

Given (i) the central role of *Dbn1*/drebrin in the LRRK2 synaptic network (Fig. 2-3), (ii) its key function as an actin regulatory protein at the synapse and (iii) its expression highly enriched at dendritic spines (Fig. S2A), we assessed drebrin protein levels at the 3 stages of life in whole brain lysates. In agreement with the qPCR results, drebrin levels are significantly lower in 1 month-old *Lrrk2* KO mice but not in older animals (Fig. 5G), further linking altered spine maturation with improper actin-dynamics in the absence of LRRK2.

Taken together, our results indicate that LRRK2 influences dendritic spine structural dynamics during brain development.

### LRRK2 knockout in human iPSC-derived neurons affects maturation and BDNF-dependent regulation of spontaneous synaptic activity

To further explore the connection between LRRK2 and BDNF activity in a human disease-relevant model, we differentiated human induced pluripotent stem cells (hiPSCs) into cortical neurons following established protocols.^70^ Wild-type and isogenic LRRK2 knockout (KO) hiPSCs^71^ were differentiated in parallel for 50, 70 and 90 days (three independent culture rounds per genotype), prior to subjecting them to patch clamp to assess spontaneous activity in the absence or presence of acute exposure to BDNF (24 hrs, 50 ng/ml) (Fig. 6A).

**Figure 6.**
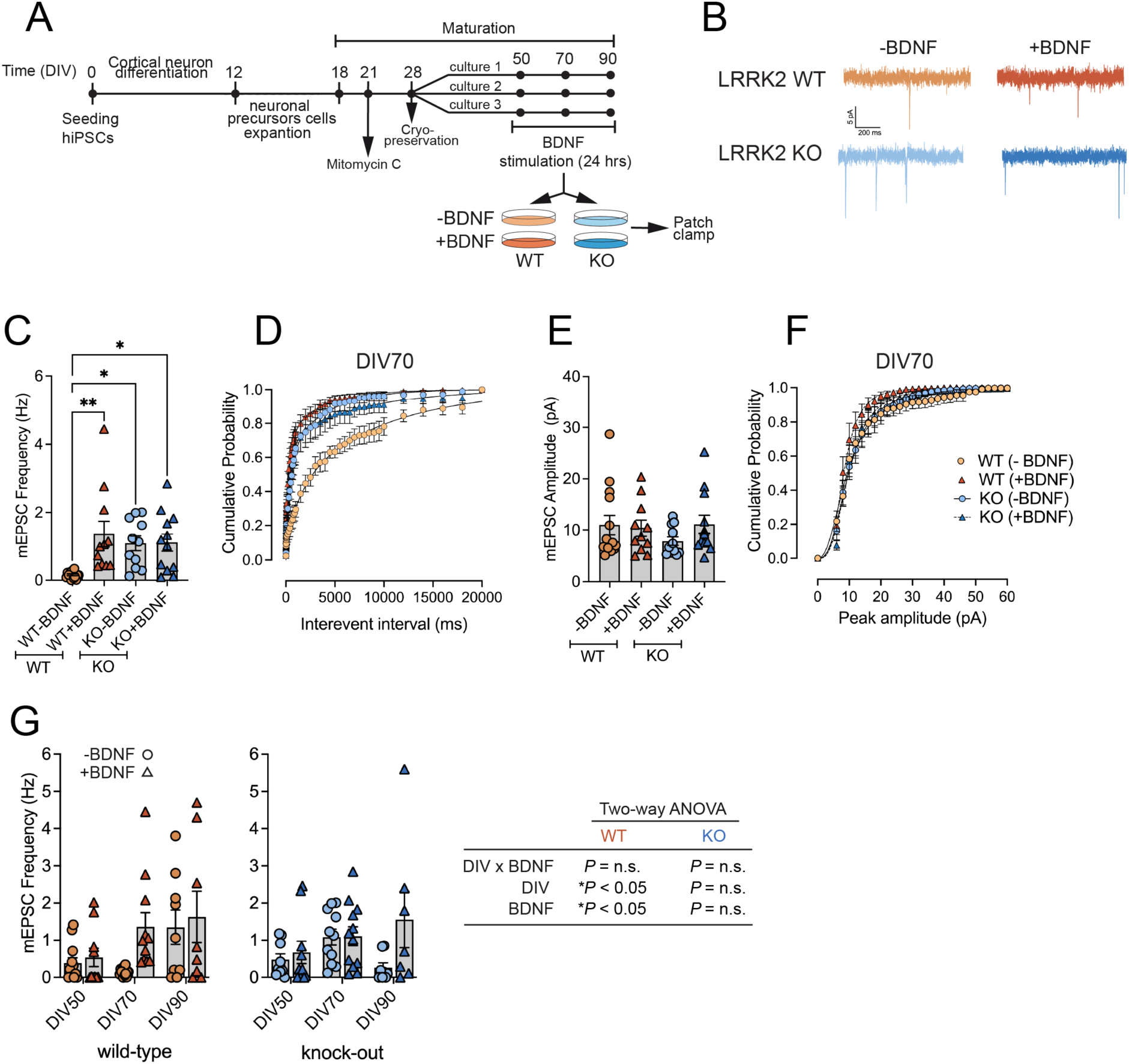
Effect of BDNF exposure on spontaneous electrical activity recorded in DIV70 LRRK2 WT and KO cortical neurons. (**A**) Schematic representation of the experimental setup. (**B**) Representative spontaneous traces of hiPSC-derived WT and KO neurons in the presence or absence of 50 ng/ml BDNF for 24 hrs. **(C)** Frequency of miniature postsynaptic excitatory currents (mEPSC) in LRRK2 WT and KO cultures (WT vehicle: 0.16 ± 0.03 Hz (*n*=15, *N*=3); WT BDNF: 1.36 ± 0.38 Hz (*n*=11, *N*=3); KO vehicle: 1.09 ± 0.21 Hz (*n*=11, *N*=3); KO BDNF: 1.11 ± 0.25 Hz (*n*=12, *N*=3). Statistical significance was determined using one-way ANOVA (WT vehicle vs. WT BDNF \*\**P*<0.01; WT BDNF vs. KO BDNF \**P*<0.05; WT vehicle vs. KO BDNF \**P*<0.05; *P*>0.05 for all the other comparisons). **(D)** Cumulative probability curves of interevent interval (IEIs) in DIV70 WT and KO -/+ BDNF treated cultures. **(E)** Amplitude of miniature postsynaptic excitatory currents (mEPSC) in DIV70 LRRK2 WT and KO cultures. BDNF treatment has no effect on the average peak amplitude (WT vehicle: 11.00 ± 1.85 pA (*n*=14, *N*=3); WT BDNF: 7.9 ± 0.84 pA (*n*=11, *N*=3); KO vehicle 10.42 ± 1.53 pA (*n*=11, *N*=3); KO BDNF 11.12 ± 1.76 pA (*n*=12, *N*=3)). Statistical significance was determined using one-way ANOVA (*P*>0.05 for all comparisons). **(F)** Cumulative probability curves of peak interval in DIV70 WT and KO -/+ BDNF treated cultures. **(G)** mEPSC frequency in WT and KO neurons across developmental stages. mEPSC frequency was measured in WT (left) and LRRK2 KO (right) neurons at DIV50, DIV70 (data plotted in C) and DIV90, in the absence (circles) or presence (triangles) of BDNF. Bars represent mean ± SEM and symbols indicate individual recorded cells. Statistical analysis was performed using two-way ANOVA (DIV × treatment) separately for WT and KO neurons. In WT neurons, significant main effects of DIV (F(2,59) = 4.61, *P* = 0.0138) and BDNF treatment (F(1,59) = 4.07, *P* = 0.0482) were detected, while the DIV × treatment interaction was not significant (F(2,59) = 1.69, *P* = 0.1936). In LRRK2 KO neurons, neither DIV (F(2,51) = 1.51, *P* = 0.2299) nor BDNF treatment (F(1,51) = 3.70, *P* = 0.0600) reached statistical significance, and no DIV × treatment interaction was observed (F(2,51) = 2.10, *P* = 0.1325).

Acute BDNF treatment led to a significant increase in the mean frequency of miniature post-synaptic excitatory currents (mEPSC) in WT cultures (Fig. 6B-C). This effect was further supported by a leftward shift in cumulative probability curves of interevent intervals (IEIs) in BDNF-treated WT neurons compared with untreated controls (Fig. 6D). In contrast, LRRK2 KO neurons displayed elevated mEPSC frequency and did not show a significant response to BDNF treatment (Fig. 6C-D). Analysis of mEPSC peak amplitudes revealed no significant differences between genotypes or treatments, as reflected by comparable mean amplitudes and overlapping cumulative amplitude distributions (Fig. 6E-F), indicating that the observed effects primarily affect synaptic event frequency rather than postsynaptic receptor function. Consistently, two-way ANOVA performed across different developmental stages (DIV50, DIV70, DIV90) revealed significant main effects of DIV and BDNF treatment in WT neurons, whereas no significant effects of DIV, BDNF, or their interaction were detected in LRRK2 KO neurons (Fig. 6G). Together, these results indicate that synaptic activity in WT neurons is regulated by both neuronal maturation and BDNF signaling, whereas LRRK2 deficiency alters this regulation. The selective effect on mEPSC frequency, but not amplitude, at DIV70 suggests that BDNF predominantly modulates presynaptic function in WT neurons. Consistent with this interpretation, quantification of presynaptic Bassoon puncta colocalized with postsynaptic Homer revealed no increase in synapse number following BDNF treatment in either genotype (Fig. S7C–D), supporting a model in which BDNF enhances neurotransmitter release probability rather than synapse formation.

## Discussion

Early synaptic alterations are thought to contribute to the pathogenesis of PD,^72^ and understanding these early events is crucial to designing effective therapeutic approaches. Previous work from our laboratories and other groups has shown that the PD-associated kinase LRRK2 regulates important synaptic processes.^73–75^ At the presynapse, LRRK2 controls the synaptic vesicle cycle through association with its WD40 domain^26,27,76,77^ and phosphorylation of a panel of presynaptic proteins, including N-ethylmaleimide sensitive fusion (NSF),^29^ synapsin I,^30^ auxilin,^33^ synaptojanin^32^ and EndophylinA.^31^ At the postsynapse, LRRK2 modulates the function of excitatory synapses in the striatum.^36,37,68,69,74,78^ However, the mechanisms by which LRRK2 affects synaptic function are still poorly elucidated. In the present study, we show that LRRK2 affects synaptic function in response to BDNF neurotrophic stimulation, promoting its relocalization to the actin-cytoskeleton in a phosphorylation-dependent manner. These findings support a model in which neurotrophic signaling engages LRRK2 to regulate cytoskeletal dynamics and synaptic function, shedding lights into early synaptic dysfunction in PD.

Our study uncovers new insights into LRRK2 regulation by neurotrophic signaling, specifically identifying BDNF as an extracellular stimulus that transiently enhances LRRK2 activity. Lrrk2 knockout neurons and differentiated SH-SY5Y LRRK2 KO cells show decreased activation of downstream AKT and ERK1/2, suggesting LRRK2 functions downstream of TrkB/BDNF and upstream of ERK1/2 and AKT. We observed high Ser935 phosphorylation under basal conditions, in agreement with mass spectrometry determination showing nearly complete phosphorylation of this residue in the whole brain.^47^ However, RAB substrates and BDNF-responsive kinases like ERK and AKT remain minimally phosphorylated at baseline, indicating they stay responsive. BDNF exposure leads to reduced Ser935 phosphorylation post-stimulation, possibly due to negative feedback mechanisms, while RAB phosphorylation did not drop below pre-stimulation levels. These findings highlight the nuanced regulation of LRRK2 activity and indicate a positive correlation between Ser935 phosphorylation and LRRK2 activity under BDNF signalling, similar to previous observations on microglia cells upon inflammatory stimuli.^79^

Moreover, using a literature-based network analysis of LRRK2 PPIs, combined with AP-MS/MS in SH-SY5Y cells and phosphoproteomics of hyperactive Lrrk2 G2019S from brain striatum, we identified a highly interconnected cluster of actin and synaptic proteins linked to LRRK2. Both BDNF-activated LRRK2 and hyperactive LRRK2 G2019S mutant are associated with GO terms related to synaptic function and cytoskeletal organization. The strong enrichment of synaptic components in the phosphoproteomic study is particularly notable given that the experiment was performed on total striatal lysates, which is a heterogeneous mixture of several cell types. This strengthens the importance of LRRK2-mediated signaling within the synaptic compartment.

The central role of synaptic dysfunction in PD neurodegeneration is increasingly recognized.^80^ The latest and largest meta-analysis of PD genome-wide association studies (GWAS), which identified 134 loci as risk factors for PD further supports this notion.^10^ The open reading frames closest to these risk signals showed strong enrichment for neuronal genes involved in dendritic spine and synapse biology (Fig. S8 and ^10^). Even though these data require functional validation (GWAS identifies risk loci rather than causal genes), they provide a genetic rationale for synaptic dysfunction as a key contributor to PD pathophysiology.^81^

Given the strong connection of LRRK2 with PD,^82,83^ elucidating the role of LRRK2 in synaptic biology will enhance our molecular understanding of how synaptic dysfunction contributes to PD.^72^ Consistent with a synaptic role for LRRK2, we found that knockout of mouse *Lrrk2* resulted in blunted BDNF-induced synapse maturation *in vitro*. Moreover, BDNF-dependent enhancement of synaptic activity is detectable in WT human iPSC-derived cortical neurons but not in KO cells. Overall, these findings suggest a mechanism by which BDNF signaling reshapes the LRRK2 interactome to promote its incorporation into a macromolecular complex required for coordinating actin dynamics during synaptic structural plasticity.

BDNF plays a key role to reorganize the actin cytoskeleton during long term potentiation (LTP) via Rho GTPases Rac1 and Cdc42.^84^ LRRK2 was previously reported to interact with Rac1^85,86^ and with other actin-related proteins.^56^ Our study shows that LRRK2 associates with the actin cytoskeleton in dendritic spines, which increases upon BDNF stimulation. We also validated the BDNF-dependency of the interaction between LRRK2 and drebrin, an actin-binding protein that stabilizes F-actin in dendritic spines.^51^ Evidence for a LRRK2-drebrin link includes increased phosphorylation at Ser339 in G2019S knockin striata and reduced drebrin levels in Lrrk2 knockout brains during development, correlating with morphological changes in dendritic protrusions. Ser339, found in the N-terminal region of human drebrin and which we identified as important for the interaction with LRRK2, lies just before the proline-rich (PP) domain of the protein. The N-terminal region of drebrin is located between the actin-binding domains and the C-terminal Homer-binding sequences, and it plays a key role in protein-protein interactions and cytoskeletal regulation.^87^ Notably, this region has been shown to interact with adafin (ADFN), a protein involved in cytoskeletal-related processes crucial for synapse formation and function. Adafin regulates puncta adherentia junctions, presynaptic differentiation, and cadherin complex assembly, all of which are essential for hippocampal excitatory synapses, spine formation, and learning and memory processes.^88^ Of note, adafin is listed among LRRK2-interacting proteins (https://www.ebi.ac.uk/intact/home), supporting a potential functional connection between LRRK2-mediated drebrin phosphorylation and the formation of adafin-drebrin complex.

It has been shown that drebrin knockout results in delayed synaptogenesis and inhibition of postsynaptic PSD95 accumulation, indicating its key role in spine maturation.^51,89,90^ During LTP, Ca^2+^ entry through N-methyl-d-aspartate (NMDA) receptors causes drebrin exodus from the spine compartment. This allows actin monomers to enter and polymerize, ultimately increasing the size of the cytoskeleton.^91^ We postulate that LRRK2 may promote drebrin exodus at the shaft of the dendrite during plasticity processes, consistent with our previous work showing that LRRK2 regulates a PKA translocation from the shaft to dendritic spines.^92^ Actin remodeling during spine enlargement is also coordinated by the ARP2/3 complex, which is activated by binding to WAVE family proteins downstream of active Rac1.^93^ Strikingly, we found that BDNF stimulates LRRK2 interaction with three out of seven subunits of the ARP2/3 complex (ACTR2, ACTR3, and ARPC2), and LRRK2 was reported to interact with WAVE proteins.^94^ Altogether these findings, in agreement with our earlier study,^38^ highlight a crucial role for LRRK2 in actin-based dendritic spine remodeling. Clearly, future studies should focus on clarifying the precise molecular dynamics and kinetics of these processes.

Actin dynamics underlie key developmental processes. In this context, our study uncovers a developmental phenotype in which loss of Lrrk2 leads to defects in dendritic protrusions morphogenesis. Interestingly, this developmental alteration is rescued with age. This is not surprising considering that the homologous kinase LRRK1 may compensate for LRRK2 deficiency, as suggested by the overt neurodegeneration observed in double *Lrrk1/Lrrk2* KO mice but absent in the single KO^95–97^ and the redundant but not overlapping pattern of expression of the two kinases.^98^ Indeed, striatal Lrrk2 expression in the rodent brain increases up to postnatal week 4,^38,67,98,99^ while the Lrrk1 transcript in this region remains relatively stable during development.^67^

Several limitations of this study should be acknowledged. First, biochemical analyses based on western blotting revealed variability in protein abundance and phosphorylation levels across experiments, likely reflecting both biological heterogeneity and technical constraints inherent to bulk tissue and cell-based assays. Second, although our electrophysiological analyses revealed genotype and BDNF-dependent effects, these measurements were performed on a limited number of independent cultures and focused on cortical neurons. Future studies using larger datasets and additional neuronal cell types will be necessary to validate and extend these findings. In particular, the study of dopaminergic neurons, the population most vulnerable in Parkinson’s disease is deemed critical important, especially considering the recent findings highlighting subtype-specific vulnerability of dopaminergic neurons in PD to account for high heterogeneity even within dopamine neurons.^72,75,100^

Several lines of evidence indicate that genetic, biochemical and cellular alterations of BDNF/TrkB signaling are linked to PD. At striatal synapses, BDNF/TrkB signaling enhances both dopamine and glutamate release and ERK1/2 activation.^101,102^ BDNF is required for the establishment of the proper number and for the survival of dopaminergic neurons in the SNpc,^103,104^ and presynaptic dopamine release in the striatum is enhanced by BDNF.^40^ BDNF levels are lower in peripheral tissues, brain, and blood of sporadic PD patients.^105^ Moreover, the V66M polymorphism modifies the risk of sporadic and mutant LRRK2-associated PD.^106,107^ Decreased BDNF concentration in serum and brain correlates with an increased degeneration of dopaminergic neurons in PD^108^ and with loss of striatal DA transporter (DAT) in patients with striatal dopaminergic neurodegeneration.^109^ Future studies should investigate whether BDNF signaling is altered in G2019S mice and hiPSCs-derived neurons with the G2019S mutation.

At the onset of PD, loss of dopaminergic axonal terminals exceeds the loss of cell bodies,^4^ implying that early synapse deterioration may be trigger axonal degeneration and, ultimately, neuronal death.^110^ In contrast to neuronal loss, which is irreversible, disease-associated synaptic dysfunctions could be rescued through the formation of new axon.^5^ Thus, focusing on early, prodromal dysfunction in PD appears key to designing effective therapeutic and preventive strategies, and our study uncovered LRRK2 as a promising disease modifying target of early-stage PD.

## Materials and methods

### Mouse strains

C57Bl/6J *Lrrk2* knock-out (KO) and G2019S-Lrrk2 knock-in mice were provided by Dr. Heather Melrose (Mayo Clinics, Florida, USA). C57B/6J or *Dbn* knock-out (KO) were previously described.^111^ Housing and handling of mice were done in compliance with national guidelines. All animal procedures were approved by the Ethical Committee of the University of Padova and the Italian Ministry of Health (license 1041/2016-PR and 105/2019), by the ‘Landesamt für Gesundheit und Soziales’ (LaGeSo; Regional Office for Health and Social Affairs in Berlin) under the permit number T-CH0025/23, and followed the guidelines approved by the Northwestern University Animal Care and Use Committee. Approximately equal numbers of males and females were used for every experiment.

### Cell cultures

#### Generation of LRRK2 KO SH-SY5Y CRISPR/Cas9 edited monoclonal cell line

The KO of LRRK2 was performed using CRISPR/Cas9-mediated genome editing technology following the protocol by Sharma *et al*.^112^ Two sgRNAs were selected among those designed by the laboratory of Zhang F (https://www.genscript.com/gRNA-detail/120892/LRRK2-CRISPR-guide-RNA.html) and one was designed using the online platform Benchling (https://www.benchling.com). All the gRNAs were synthetized by Sigma-Aldrich. The oligos pairs encoding the 20-nt guide sequence were annealed and ligated into the pSpCas9(BB)-2A-Puro (PX459) V2.0 vector (Addgene, Watertown, MA, US) and then amplified in chemically competent E. coli StbI3 cells (ThermoFisher ScientificTM). Human neuroblastoma-derived SH-SY5Y cells were transfected using Lipofectamine 2000 (Invitrogen) and subjected to puromycin selection. The selected cells were diluted to obtain monoclonal cell lines. Approximately one week after plating, the colonies were inspected for a clonal appearance and progressively expanded. Finally, the deletion of LRRK2 was verified in multiple lines by western blot analysis with LRRK2 specific antibody (Fig. S5).

#### SH-SY5Y cell line maintenance, differentiation and treatments

SH-SY5Y cells (naïve, LRRK2 KO, expressing GFP or GFP-LRRK2 wild type) were cultured in a mixture (1:1) of Dulbecco’s Modified Eagle’s Medium (DMEM, Biowest) and Ham’s F-12 Nutrient Mixture (F12, Biowest), supplemented with 10% (v/v) Fetal Bovine Serum (FBS, Thermo Fisher Scientific) and 1% Penicillin/Streptomycin solution (PS, GIBCO Life Technologies).

To promote N-type (neuronal-like cells) cell differentiation, cells were plated in DMEM/F12 containing 1% PS, 1% FBS and 10μM of all-trans-retinoic acid (RA, Sigma-Aldrich). At regular intervals of 48 hours, RA was newly provided to the medium. Cells were differentiated for 6 days and then subjected to the treatments.

SH-SY5Y cells were starved for 5 hours in DMEM/F12 supplemented with 1% PS and then stimulated with 100ng/mL BDNF (50240-MNAS, Sino Biological) for different time periods (5, 15, 30 and 60 minutes) in serum-free medium. LRRK2 kinase activity was inhibited pre-treating cells with 0.5μM MLi-2 (ab254528, Abcam) for 90 minutes. For the detection of phospho-S935 and phospho-Rabs, 80 µg of total protein were loaded, whereas 30 µg were used for the detection of phospho-AKT and phospho-ERK1/2, as well as for total proteins.

### HEK293T cell line maintenance and transfection

HEK293T cells (Life technologies) were cultured in DMEM (Biowest) supplemented with 10% (v/v) FBS (Thermo Fisher Scientific) and 1% PS (GIBCO Life Technologies). Cells were plated in 100mm dishes and, once they reached 80% confluency, they were transfected with polyethyleneimine (PEI, Polysciences) at 1:2 DNA to PEI ratio (v/w). Flag-LRRK2 was co-expressed with YFP-drebrin full-length, N-terminal domain (1-256 aa), or C-terminal domain (256-649 aa).

### Primary mouse cortical neuron preparation, transfection and treatments

Primary cortical neurons from *Lrrk2* WT and KO C57BL/6J mice were derived from postnatal mouse (P0) exploiting the Papain Dissociation System (Worthington Biochemical Corporation). Cortices were incubated in Papain solution (Papain and DNase solution in Earle’s Balanced Salt Solution) for 40 minutes at 37°C. Subsequently, tissue was triturated and centrifugated for 5 minutes at 200g. The supernatant was discarded, and the pellet was resuspended in Stop solution [DNase solution and Trypsin inhibitor solution (15,5mg/mL) in Earle’s Balanced Salt Solution]. For 10 minutes the tissue was allowed to precipitate and then the supernatant was pipetted drop-by-drop on 5mL of 10/10 solution (10 µg/mL Trypsin inhibitor and 10 µg/mL BSA in Earle’s Balanced Salt Solution) and centrifugated for 10 minutes at 100g. The pellet was resuspended in Neurobasal A medium (GIBCO Life Technologies) supplemented with 2% B27 supplement (GIBCO Life Technologies), 0.5 mM L-glutamine (GIBCO Life Technologies), 100 units/mL penicillin, and 100 μg/mL streptomycin (GIBCO Life Technologies). Cells were diluted and counted in 0.4% Trypan blue. Neurons were plated at 1000-1500 cells/mm^2^ onto 6-well plates or at 200 cells/mm^2^ on 12mm glass coverslips in 24-well plates and maintained in culture for 14 days prior to the treatments. After 7 days, 50% of the Neurobasal medium was removed and replaced with fresh one.

At DIV14, *Lrrk2* WT and KO mature neurons cultured in the 6-well plates for western blot analysis were treated with MLi-2 (0.5μM) for 90 minutes and with BDNF (100ng/mL) for 5, 30, 60 and 180 minutes in Neurobasal completed medium. At DIV4, Lrrk2 WT and KO primary neurons plated in 24-well plates were transfected with lipofectamine (Lipofectamine 2000, Invitrogen) in a 1:2 ratio with DNA (v/w). The transfection was carried in Opti-MEM (GIBCO Life Technologies) for 45 minutes. Transfected neurons were then maintained in culture for additional 9-10 days. At DIV14, neurons were treated for 24 hours with 100ng/mL BDNF to study the spine maturation process.

### Media composition for cortical neuron differentiation from hiPSC

Neuronal maintenance media (NMM): 1:1 mixture of N2 media (DMEM/F12 GlutaMax supplemented with 1X N2, 50 units/mg/ml Penicillin/Streptomycin solution, 1X MEM Non-Essential Amino Acids Solution (100X), 100µM 2-Mercaptoethanol (ThermoFisher Scientific), 1mM sodium pyruvate and 5 µg/ml insulin (Merck)) and B27 media (Neurobasal supplemented with 1X B27 and 2mM L-Glutamine (ThermoFisher Scientific). Neuronal induction media (NIM): NMM media supplemented with 100nM LDN193189 (Sigma Aldrich) and 10µM SB431542 (Bio-Techne).

### hiPSC cell culture, cortical neuron differentiation and treatments

Isogenic human induced-pluripotent stem cell lines (hiPSC) of WT and LRRK2 KO, made and kindly provided by Mark R Cookson’s laboratory^71^ were cultured using E8 media (Thermo Fisher Scientific) in a 6-well plate coated with Cultrex. Cortical neuronal differentiation was carried out according to Shi, *et al*.^70^ Briefly, upon reaching 100% confluency, media was changed to neuronal induction media (NIM) and is marked as 0 days *in vitro* (DIV0). Daily NIM media change was carried out till DIV12, upon which neuronal precursor cells (NPCs) are generated. On DIV12, NPCs were expanded via passage at 1:2 ratio, supplemented with neuronal maintenance media (NMM) media and 20 ng/ml FGF2 (Bio-Techne). Neurogenesis was initiated on DIV18 with the removal of FGF2 and the cells supplemented with NMM only changing every 2 days. At DIV25-30, neurons were cryopreserved in CryoStor CS10 solution (Stem Cell technologies).

For electrophysiological recordings and immunocytochemistry/immunofluorescence (ICC/IF), thawed neurons were plated on Cultrex coated 10mm coverslips at a density of 60k/cm^2^. All cells were treated with 1 ug/ml Mitomycin C (Sigma-Aldrich) for 1 hour at 37°C, 48-72 hours after plating to remove any proliferating cells. Neurons were maintained in NMM until used for experiments. BDNF treatment was carried out with cells incubated with 50 ng/ml BDNF (Bio-Techne) for 24 hours prior to use in electrophysiological recordings and ICC/IF.

### Electrophysiological recordings

Whole-cell patch-clamp recordings were performed on cortical neurons at DIV70-75 (mentioned as DIV70 from here on) in voltage clamp at Vh −70 mV and the membrane test function was used to determine intrinsic membrane properties ∼1 min after obtaining whole-cell configuration, as described previously.^113^ Briefly, coverslip containing neurons were transferred to warner bath (RC-26G) and maintained at 35 ± 2 °C through constant perfusion using in-line heater and controller (Warner Instruments) with extra cellular solution (ECS) containing (in mM unless stated): 145 NaCl, 3 KCl, 2 MgCl_2_, 2 CaCl_2_, 10 glucose, 10 HEPES, pH 7.4, 310 mOsm. Tetrodotoxin (TTX 0.2μM), and picrotoxin (PTX 100μM), were added before use to block sodium and GABA_A_ currents. Pipette resistance (Rp) of glass electrodes was 4–6 MΩ when filled with (in mM): 130 Csmethanesulfonate, 5 CsCl, 4 NaCl, 2 MgCl2, 5 EGTA, 10 HEPES, 5 QX-314, 0.5 GTP, 10 Na2-phosphocreatine, and 5 MgATP, 0.1 spermine, pH 7.2, 300 mOsm. Data was acquired by Multiclamp700B amplifier and digidata 1550B and signals were filtered at 2kHz, digitized at 10 kHz, and analyzed in Clampfit 10 (MolecularDevices). Tolerance for series resistance (Rs) was <35 MΩ and uncompensated; ΔRs tolerance cut-off was <20%. mEPSCs were analyzed using Clampfit10 (threshold 5pA); all events were checked by eye. Data are presented as mean ± s.e.m. where n is cells from a minimum of 3 separate cultures (culture N in brackets).

### Cells and tissues lysis, SDS-PAGE and Western blotting analysis

Immortalized cells, neurons and mouse brain tissues were lysed for 30 minutes on ice in appropriate volume of cold RIPA lysis buffer (Cell Signaling Technologies) supplemented with protease inhibitor cocktail. Protein concentration was assessed by performing BCA assay (Thermo Fisher Scientific). Proteins were solubilized in sample buffer (200mM Tris-HCl pH 6.8, 8% SDS, 400mM DTT, 40% glycerol, q.s. Bromophenol Blue) and resolved by SDS-PAGE on 8% Tris polyacrylamide homemade gels in Tris-glycine-SDS running buffer for the AP-MS experiments, and on 4-20% polyacrylamide gels (GenScript® Express Plus PAGE) in Tris-MOPS-SDS running buffer (GenScript® Running Buffer Powder) for the other cases. Proteins were transferred on PVDF (polyvinylidene difluoride, Bio-Rad) membranes using a semi-dry transfer system (Trans-Blot® Turbo Transfer System, Bio-Rad). Non-specific binding sites were blocked incubating membranes with 5% non-fat dry milk diluted in 0.1% Tween-20 Tris-buffered saline (TBS-T) for 1 hour at room temperature under agitation. Membranes were subsequently incubated overnight at 4°C with primary antibodies in 5% non-fat dry milk in TBS-T or in 5% BSA in TBS-T. After three washes in TBS-T at room temperature, membranes were incubated for 1 hour at room temperature with horseradish peroxidase (HRP)-conjugated goat anti-mouse or anti-rabbit IgG. After three more washes in TBS-T, proteins were visualized using chemiluminescence (Immobilon ECL western HRP substrate, Millipore). Densitometric analysis was carried out using the Image J software. The antibodies used for western blotting are as follows: rabbit α-LRRK2 (MJFF2 c41-2, ab133474, Abcam, 1:300); rabbit α-phospho-Ser935 LRRK2 (ab133450, Abcam, 1:300); mouse α-phospho-Ser473-AKT (sc-293125, Santa Cruz Biotechnology, 1:500); rabbit α-AKT (9272S, Cell Signaling Technology, 1:1000); rabbit α-phospho [(Thr202/Tyr204, Thr185/Tyr187)-ERK1/2 (12-302, Millipore, 1:1500); rabbit α-ERK1/2 (4695, Cell Signaling Technology, 1:1000); mouse α-GAPDH (CSB-MA000195, 1:5000); mouse α-βIII tubulin (T8578, Sigma-Aldrich, 1:40000); mouse α-DREBRIN (MA1-20377, Thermo Fisher Scientific, 1:500); α-Flag M2-HRP (A8592, Sigma-Aldrich, 1:5000); α-mouse IgG-HRP (A9044, Sigma-Aldrich, 1:80000); α-rabbit IgG-HRP (A9169, Sigma-Aldrich, 1:16000).

### Staining on mammalian cells and brain tissue

#### Immunocytochemistry and confocal microscopy of primary neurons

Mouse primary cortical neurons derived from *Lrrk2* WT C57BL/6 and KO pups (P0) were fixed using 4% paraformaldehyde (PFA, Sigma-Aldrich) in PBS1X pH 7.4 for 20 minutes at room temperature and, after three washes in PBS1X, they were subjected to staining protocol. Cells were firstly permeabilized with 0.3% Triton-X in PBS1X for 5 minutes and then saturated in blocking buffer [1% Bovine serum Albumin (BSA) Fraction V, 0.1% Triton-X, 50mM Glycine, 2% goat serum in PBS1X] for 1 hour at room temperature in agitation. The primary and secondary antibodies incubation steps were carried out in working solution (20% blocking buffer in PBS1X) for 1 hour at room temperature. Both incubations were followed by three washes in working solution. Nuclei staining was performed in Hoechst 33258 (Invitrogen, 1:10000 dilution in PBS1X) for 5 minutes and followed by three rinses in PBS1X. After been cleaning in distilled water, coverslips were mounted on microscope slides with Mowiol® mounting medium. Immunofluorescence z-stack images (z-stack thickness: 0.5 μm x 6 = 3 μm) were obtained on Zeiss LSM700 laser scanning confocal microscope exploiting a 63X oil immersion objective.

The antibodies used for immunocytochemistry are as follows: mouse α-PSD95 (ab2723, Abcam, 1:200), rabbit α-MAP2 (sc-20172, Santa Cruz, 1:200); mouse α-drebrin (MA1-20377, Thermo Fisher Scientific, 1:400); goat anti-mouse-Alexa Fluor 568 (A11004, Invitrogen), goat anti-mouse-Alexa Fluor (A11004, Invitrogen), goat anti-rabbit-Alexa Fluor 488 (A11034, Invitrogen), rabbit-Alexa Fluor 568 (A11036, Invitrogen), goat anti-rabbit-Alexa Fluor 405.

#### Immunostaining and Image Analysis of hiPCS-derived cortical neurons

For immunocytochemistry DIV70 cells on coverslips were fixed in 4% paraformaldehyde (PFA) for 10 minutes. Cells were then permeabilized and blocked using 0.4% Tween with 10% normal goat serum (NGS), in PBS1X for 1 hour. Primary antibodies were incubated overnight at 4°C in PBST plus 10% NGS. Cells were washed 3× with PBST before 1 hour incubation at room temperature with secondary antibodies (α-guinea pig Alexa-488, α-rabbit Alexa 568, α-chicken 647 along with DAPI) from Molecular probes and Jackson Laboratories). Primary antibodies were chicken anti-microtubule associated protein 2 (MAP2) (Antibodies A85363, dil. 1:1000), rabbit anti-homer 1 (Synaptic Systems 160003, dil. 1:500), and guinea pig anti-bassoon (Synaptic Systems 141004, dil. 1:500). Coverslips were slide mounted with FluorSave (Sigma-Aldrich) and all images were acquired on Leica confocal microscope as 0.45 μm z-stacks at 60× magnification (flattened with the max projection function for cluster analysis). Single 60× images were used for analyses. Excitation and acquisition parameters were constrained across all experimental paired (culture) acquisitions. All images were analyzed using IMARIS microscopy image analysis software (v10.0.0 Oxford Instruments). Data are presented as mean ± s.e.m.

#### Golgi-Cox staining

Animals were terminally anesthetized and transcardially perfused with 0.9% saline. Half of each brain was incubated with Golgi-Cox solution (Potassium dichromate, Mercuric chloride, Potassium chromate prepared according to Zaqout *et al.*^114^ in the dark at room temperature for 14 days and then transferred in 30% sucrose solution in PBS1X. Brains were cut with a vibratome in 100 μm thick slices. The sections were then blotted by pressing an absorbent paper moistened with sucrose solution onto the slides and dried for 7-10 minutes. The samples were then subjected to color development procedure consisting in the following steps: 1. two 5-minutes washes in distilled water; 2. 5-minutes dehydration step in 50% ethanol; 3. 10-minutes incubation step in 20% ammonium hydroxide; 4. 5-minutes wash in distilled H_2_O; 5. 8-minutes incubation step in 5% sodium thiosulfate at room temperature in the dark; 6. two additional 1-minute rinses in distilled H_2_O; 7. dehydration in ascending grades of ethanol (70%, 95% and 100%). After two final 6-minutes incubations with xylene, the slides were covered with Eukitt® mounting medium. Z-stack images (z-stack thickness between 0.5 μm x 30 = 15 μm and 0.5 μm x 80 = 40 μm) were acquired with Zeiss LSM700 laser scanning confocal microscope using 100X/1,40 Oil DIC M27 immersion objective with phase contrast acquisition mode. All images were analyzed using freely available RECONSTRUCT software and according to Risher *et al*.^115^ 3-4 neurons per mice (n=3 per genotype) were acquired in the dorsal striatum and n≌4 segments per neurons were analyzed.

#### Electron Microscopy

Coronal brain slices were fixed with 2.5% glutaraldehyde in 0.1M sodium cacodylate pH 7.4 buffer overnight at 4° C, postfixed in 1% osmium tetroxide for 1h at 4° C, ethanol dehydrated, then infiltrated in a mixture of EMbed 812 (Electron Microscopy Sciences) epoxy resin and absolute ethanol 1:1, and then finally embedded in EMbed 812 (Electron Microscopy Sciences) epoxy resin. Ultrathin sections were obtained with a Leica EM UC7 ultramicrotome and subsequently counterstained with uranyl acetate and lead citrate. Samples were then examined with a Tecnai G2 (Fei-Thermo Fisher) transmission electron microscope operating at 120 kV, digital images were acquired using a Veleta (Olympus Soft Imaging Solutions) digital camera. Images were acquired at the level of dorsal striatum in n=4 mice per genotype (1-month old mice) or n=3 mice per genotype (18-months old mice). n≌20 synapses per mice were analyzed in 1-month old mice; n≌40 synapses per mice were analyzed in 18-months old mice. We gratefully thank the DeBio Imaging facility at the University of Padova for their support with the staining procedure.

#### Quantitative PCR

Total RNA was isolated from midbrain, striatum, and cortex of 1 month-old animals (n=6 animals per group, Lrrk2 WT and Lrrk2 KO) using Animal Tissue RNA Purification Kit (Norgen), according to manufacturer’s instruction. After extraction, RNA concentration was quantified with NanoDrop 2000C spectrophotometer (Thermo Fisher Scientific). After DNAse treatment (Thermo Fisher Scientific, according to manufacturer’s protocol), complementary DNA (cDNA) was generated using qRT SuperMix (Bimake). The cDNAs were used for quantitative PCR (qPCR) exploiting iTaq Universal SYBR® Green Supermix and CFX96 RealTime System (BioRad) for 40 cycles. All samples were run in triplicated, and transcripts levels normalized against the geometrical means of RPL27, actin, and GAPDH relative abundance (housekeeping genes). Data shown were produced using Bio-Rad CFX Manager software and analyzed according to ddCt algorithm.

Primers (5’-3’) used are:

*Bdnf* FW: GGCTGACACTTTTGAGCACGTC *Bdnf* REV: CTCCAAAGGCACTTGACTGCTG *TrkB* FW: TGAGGAGGACACAGGATGTTGA *Trkb* REV: TTCCAGTGCAAGCCAGTATCTG

*Shank3* FW: ACCTTGAGTCTGTAGATGTGGAAG *Shank3* REV: GCTTGTGTCCAACCTTCACGAC *Psd95* FW: GGTGACGACCCATCCATCTTTATC *Psd95* REV: CGGACATCCACTTCATTGACAAAC *Arpc2* FW: GAGTCACAGTAGTCTTCAGCACG *Arpc2* REV: AGGTTCCCTGTGGCTGAAAAGG *Dbn1* FW: AGAAGTCGGAGTCAGAGGTGGA *Dbn1* REV: ATGCCACTCGTTCCTGCTGTCT *Darpp32* FW: AGATTCAGTTCTCTGTGCCCG *Darpp32* REV: GGTTCTCTGATGTGGAGAGGC housekeeping genes

*Rpl27* FW: AAGCCGTCATCGTGAAGAACA *Rpl27* REV: CTTGATCTTGGATCGCTTGGC *GAPDH* FW: AGGTCGGTGTGAACGGAT TTG *GAPDH* REV: TGTAGACCATGTAGTTGAGGTCA *ACTB* FW: CAACGGCTCCGGCATGTG

*ACTB* REV: CTCTTGCTCTGGGCCTCG

#### Protein purification from mammalian cells

For the affinity purification (AP) protocol, SH-SY5Y GFP and SH-SY5Y GFP-LRRK2 cells were plated onto 100mm dishes and differentiated for 6 days. SH-SY5Y GFP-LRRK2 cells were treated with BDNF (100ng/mL) or with an equal volume of vehicle for 15 minutes, while SH-SY5Y OE-GFP cells were left untreated. Cells were harvested in lysis buffer (20mM Tris-HCl pH 7.5, 150mM NaCl, 1mM EDTA, 1% Tween 20, 2.5mM sodium pyrophosphate, 1mM β-glycerophosphate, 1mM sodium orthovanadate) supplemented with protease inhibitor cocktail (Sigma-Aldrich). Lysates were incubated end-over-end with GFP-TrapA beads (ChromoTek) overnight at 4°C. Immunocomplexes were subsequently washed ten times with buffers containing a decreasing amount of NaCl and Tween20 and incubated with sample buffer 2X. The eluted proteins were resolved by SDS-PAGE and processed for western blot analysis or Mass Spectrometry (MS).

HEK293T cells were transfected as described above and drebrin constructs were immunoprecipitated with GFP-trap resin following the same protocol used for the SH-SY5Y cells. The eluded samples were resolved by SDS-PAGE and processed for western blot analysis to visualize the co-precipidated LRRK2 using Flag antibodies.

#### LC-MS/MS and data analysis

Three biological replicates of GFP-trap purifications of SH-SY5Y cells were processed for the proteomics experiments. Gels were stained with Colloidal Coomassie Brilliant Blue (0.25% Brilliant Blue R-250, 40% ethanol, 10% acetic acid in milli-Q water) for at least 1 hour and then rinsed in destaining solution (10% isopropanol, 10% acetic acid in milli-Q water). Gel slices were cut into small pieces and subjected to reduction with dithiothreitol (DTT 10 mM in 50 mM NH4HCO3, for 1 h at 56 °C), alkylation with iodoacetamide (55 mM in 50 mM NH4HCO3, for 45 min at RT and in the dark), and in-gel digestion with sequencing grade modified trypsin (Promega, 12.5 ng/μL in 50 mM NH4HCO3). Samples were analyzed with a LTQ-Orbitrap XL mass spectrometer (Thermo Fisher Scientific) coupled to a HPLC UltiMate 3000 (Dionex – Thermo Fisher Scientific) through a nanospray interface. Peptides were separated at a flow rate of 250 nL/min using an 11-cm-long capillary column (PicoFrit, 75-μm ID, 15-μm tip, New Objective) packed in house with C18 material (Aeris Peptide 3.6 μm XB C18; Phenomenex). A linear gradient of acetonitrile/0.1% formic acid from 3 to 40% was used for peptide separation and the instrument operated in a data dependent acquisition mode with a Top4 method (one full MS scan at 60,000 resolution in the Orbitrap, followed by the acquisition in the linear ion trap of the MS/MS spectra of the four most intense ions). Two technical replicates were acquired for each biological replicate. Raw data files were analyzed using MaxQuant and Andromeda software package^116^ and searched against the human section of the UniProt database (version September 2020, 75093 entries) concatenated with a database of common contaminants found in proteomic experiments. Trypsin was selected as digesting enzyme with up to two missed cleavages allowed, carbamidomethylation of cysteine residues was set as a fixed modification and methionine oxidation as a variable modification. A search against a randomized database was used to assess the false discovery rate (FDR), and data were filtered to remove contaminants and reverse sequences and keep into account only proteins identified with at least two peptides and a FDR ≤ 0.01, both at peptide and protein level.

Intensity values were retrieved for 350 proteins. A single step of data QC was applied to the intensity values: proteins whose intensity value was 0.00 for both the 2 technical replicates in at least 1 of the 3 pull-down experiments performed with LRRK2-GFP were removed. This is a stringent QC step, but it was implemented to remove proteins with no triplicate data available for the LRRK2 pull-down as this could have occurred because of genuine low levels of pull-down (values close to the limit of detection, missing not at random values) or because of technical issues with the single sample (loss/poor quality of sample, missing at random values). QC reduced the number of proteins in the analysis from 350 to 269 entries.

Intensity data were then processed according to Aguilan *et al.*^117^ Briefly, intensity data were log2 transformed and normalized to the median of all proteins within the same experiment. Missing data (0.00 intensity values) were considered missing not at random thus they were imputed via probabilistic minimum imputation (random values within less than 2.5 standard deviations from the distribution of intensity values per the single experiment, 0.3 max variability).^117,118^

Finally, fold change (GFP-LRRK2 vs GFP and GFP-LRRK2 BDNF treated vs GFP-LRRK2 untreated) and associated p-value (two-tailed, paired t-test) were calculated and visualized.

### Phosphoproteomics analysis

#### Protein processing for MS

Striata from 8 weeks mice were dissected and rapidly homogenized in four volumes of ice-cold Buffer A (0.32 M sucrose, 5 mM HEPES, pH7.4, 1 mM MgCl_2_, 0.5 mM CaCl_2_) supplemented with Halt protease and phosphatase inhibitor cocktail (Thermo Fisher Scientific) using a Teflon homogenizer (12 strokes). Homogenized brain extract was centrifuged at 1400g for 10 minutes. Supernatant (S1) was saved and pellet (P1) was homogenized in buffer A with a Teflon homogenizer (five strokes). After centrifugation at 700 g for 10 minutes, the supernatant (S1’) was pooled with S1. Pooled S1 and S1′ were centrifuged at 13,800 g for 10 minutes to the crude synaptosomal pellet (P2). The crude synaptosomal pellet (P2) was homogenized in buffer B (0.32M sucrose, 6mM Tris, pH 8.0) supplemented with protease and phosphatase inhibitors cocktail with a Teflon homogenizer (five strokes) and was carefully loaded onto a discontinuous sucrose gradient (0.8 M/1 M/1.2 M sucrose solution in 6 mM Tris, pH 8.0) with a Pasteur pippete, followed by centrifugation in a swinging bucket rotor for 2 hours at 82,500g. The synaptic plasma membrane fraction (SPM) in the interphase between 1 M and 1.2 M sucrose fractions was collected using a syringe and transferred to clean ultracentrifuge tubes. 6 mM Tris buffer was added to each sample to adjust the sucrose concentration from 1.2 M to 0.32 M and the samples were centrifuged in a swinging bucket rotor at 200,000g for 30 minutes. The supernatant was removed and discarded. Added to the pellets the lysis solution containing 12 mM sodium deoxycholate, 12 mM sodium lauroyl sarcosinate, 10 mM TCEP, 40 mM CAA, and phosphatase inhibitor cocktail (Millipore-Sigma) in 50 mM Tris·HCl, pH 8.5, and incubated 10 min at 95°C with vigorous shaking. Sonicated the pellets several times with a sonicator probe and boiled again for 5 minutes. Centrifuged at 16,000g for 10 minutes to remove the debris and collected supernatant. The samples were diluted fivefold with 50 mM triethylammonium bicarbonate and analyzed by BCA to determine protein concentration. The samples were then normalized to 300μg protein in each and digested with 6μg Lys-C (Wako) for 3 hours at 37°C. 6μg trypsin was added for overnight digestion at 37°C. The supernatants were collected and acidified with trifluoroacetic acid (TFA) to a final concentration of 1% TFA. Ethyl acetate solution was added at 1:1 ratio to the samples. The mixture was vortexed for 2 minutes and then centrifuged at 16,000g for 2 minutes to obtain aqueous and organic phases. The organic phase (top layer) was removed, and the aqueous phase was collected, dried completely in a vacuum centrifuge, and desalted using Top-Tip C18 tips (Glygen) according to manufacturer’s instructions. The samples were dried completely in a vacuum centrifuge and subjected to phosphopeptide enrichment using PolyMAC Phosphopeptide Enrichment kit (Tymora Analytical) according to manufacturer’s instructions, and the eluted phosphopeptides dried completely in a vacuum centrifuge.

#### LC-MS/MS Analysis

The full phosphopeptide sample was dissolved in 10.5 μL of 0.05% trifluoroacetic acid with 3% (vol/vol) acetonitrile and 10 μL of each sample was injected into an Ultimate 3000 nano UHPLC system (Thermo Fisher Scientific). Peptides were captured on a 2-cm Acclaim PepMap trap column and separated on a 50-cm column packed with ReproSil Saphir 1.8 μm C18 beads (Dr. Maisch GmbH). The mobile phase buffer consisted of 0.1% formic acid in ultrapure water (buffer A) with an eluting buffer of 0.1% formic acid in 80% (vol/vol) acetonitrile (buffer B) run with a linear 90-min gradient of 6–30% buffer B at flow rate of 300 nL/min. The UHPLC was coupled online with a Q-Exactive HF-X mass spectrometer (Thermo Fisher Scientific). The mass spectrometer was operated in the data-dependent mode, in which a full-scan MS (from m/z 375 to 1,500 with the resolution of 60,000) was followed by MS/MS of the 15 most intense ions (30,000 resolution; normalized collision energy −28%; automatic gain control target (AGC) - 2E4, maximum injection time - 200 ms; 60sec exclusion].

#### LC-MS Data Processing

The raw files were searched directly against the mouse database with no redundant entries, using Byonic (Protein Metrics) and Sequest search engines loaded into Proteome Discoverer 2.3 software (Thermo Fisher Scientific). MS1 precursor mass tolerance was set at 10 ppm, and MS2 tolerance was set at 20ppm. Search criteria included a static carbamidomethylation of cysteines (+57.0214 Da), and variable modifications of phosphorylation of S, T and Y residues (+79.996 Da), oxidation (+15.9949 Da) on methionine residues and acetylation (+42.011 Da) at N terminus of proteins. Search was performed with full trypsin/P digestion and allowed a maximum of two missed cleavages on the peptides analyzed from the sequence database. The false-discovery rates of proteins and peptides were set at 0.01. All protein and peptide identifications were grouped and any redundant entries were removed. Only unique peptides and unique master proteins were reported.

#### Label-free Quantitation Analysis

All data were quantified using the label-free quantitation node of Precursor Ions Quantifier through the Proteome Discoverer v2.3 (Thermo Fisher Scientific). For the quantification of phosphoproteomic data, the intensities of phosphopeptides were extracted with initial precursor mass tolerance set at 10 ppm, minimum number of isotope peaks as 2, maximum ΔRT of isotope pattern multiplets – 0.2 min, PSM confidence FDR of 0.01, with hypothesis test of ANOVA, maximum RT shift of 5 min, pairwise ratio-based ratio calculation, and 100 as the maximum allowed fold change. For calculations of fold-change between the groups of proteins, total phosphoprotein abundance values were added together, and the ratios of these sums were used to compare proteins within different samples.

#### Bioinformatics

The LRRK2 interactome was constructed following the procedure reported in Zhao *et al.*^119^ Briefly, protein interactions reported for LRRK2 in peer-reviewed literature were considered if the interaction was replicated in at least 2 different papers or identified with 2 different interaction detection methods. The obtained LRRK2 protein interaction network was analysed in Zhao *et al.*^119^ to identify topological clusters using the FastGreed R Package (based on degree centrality). Here we report one of the identified clusters that is enriched for synaptic functions. Functional enrichment was performed using g:Profiler, g:GOSt (https://biit.cs.ut.ee/gprofiler/gost)^120^ with Gene Ontology Biological Processes (GO-BPs) and SynGO (https://www.syngoportal.org/).

#### Statistical analysis

Statistical analyses and plotting were performed with GraphPad-Prism9. Unpaired, two-tailed t-test or Shapiro-Wilk test were used for experiments comparing two groups; one-way ANOVA was used for experiments comparing three or more groups; two-way ANOVA was used when multiple groups and two factors (i.e., genotype and treatment) were to be compared. Šídák’s multiple comparisons test was used when determining statistical significance for comparisons between groups.

## Data availability

The authors confirm that the data supporting the findings of this study are available within the article and its supplementary material. Protein composition of the LRRK2 cluster enriched for synaptic functions can be obtained from Zhao *et al*.^119^

## Acknowledgments

We are very grateful to Dr. Mark R Cookson and Alexandra Beilina (NIA, NIH, USA) for providing WT and LRRK2 KO human iPSC lines used for electrophysiological measurements.

## Supplementary figures

**Figure S1.**
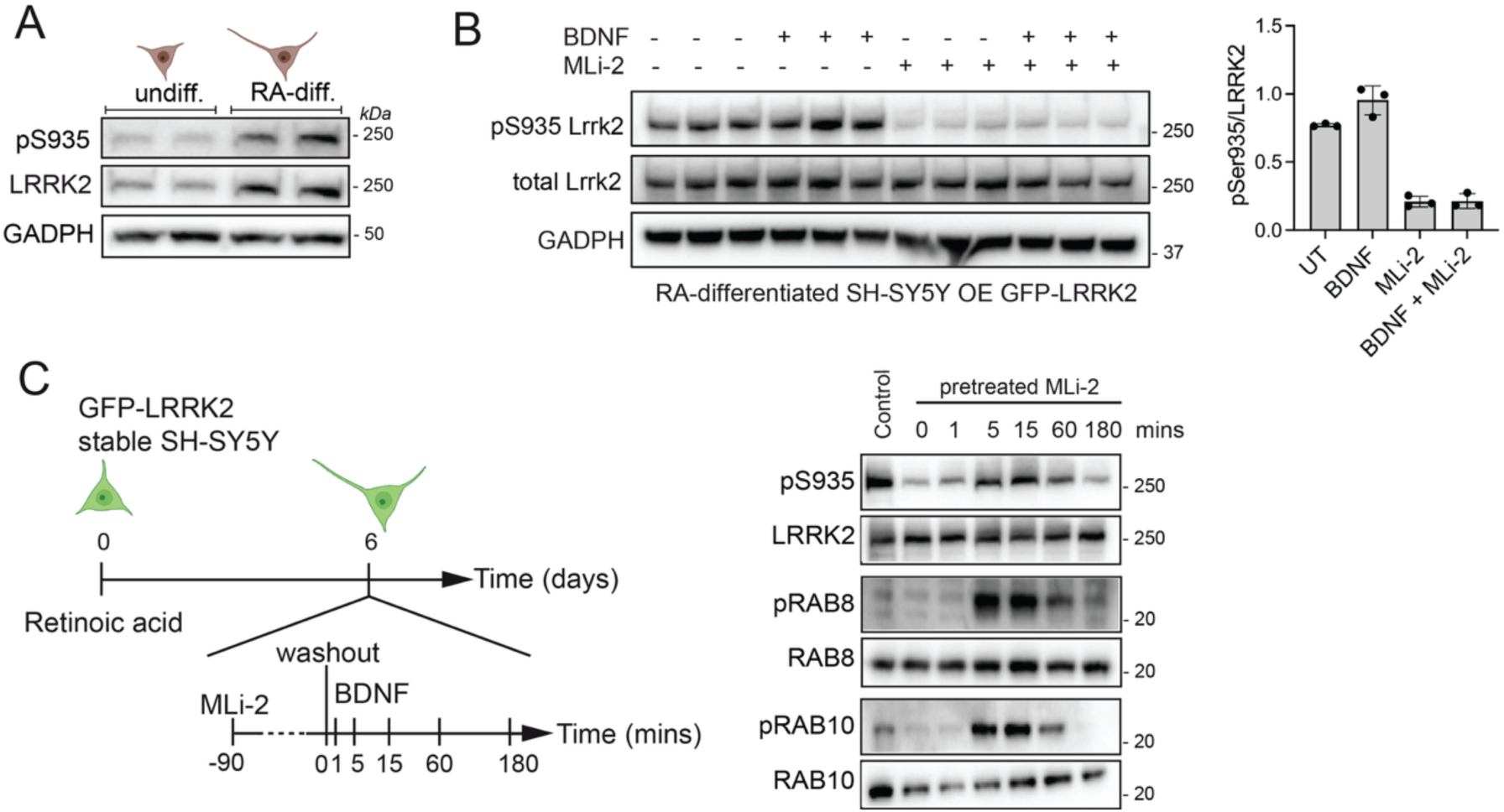
**(A)** Western blot analysis of undifferentiated versus differentiated naïve SH-SY5Y and western blot with anti-phospho-Ser935 LRRK2 and total LRRK2 antibodies. **(B)** Cells stably overexpressing GFP-LRRK2 treated with 100 ng/ml BDNF for 5 mins or with 500 nM MLi-2 for 90 mins. **(C)** On the left, schematic representation of experimental setting. On the right, BDNF stimulation of GFP-LRRK2 SH-SY5Y cells prior to MLi-2 (500 nM) treatment for 90 minutes (MLi-2 for 90’, washout, BDNF stimulation for 1’, 5’, 15’, 60’ and 180’). The timepoint 0’ correspond to non BDNF treated. Western blot were performed using anti LRRK2 (phospho-Ser935 and total), anti-RAB10 (phospho Thr73 and total) and anti-RAB8 (pan-phopho-RABs and total RAB8).

**Figure S2.**
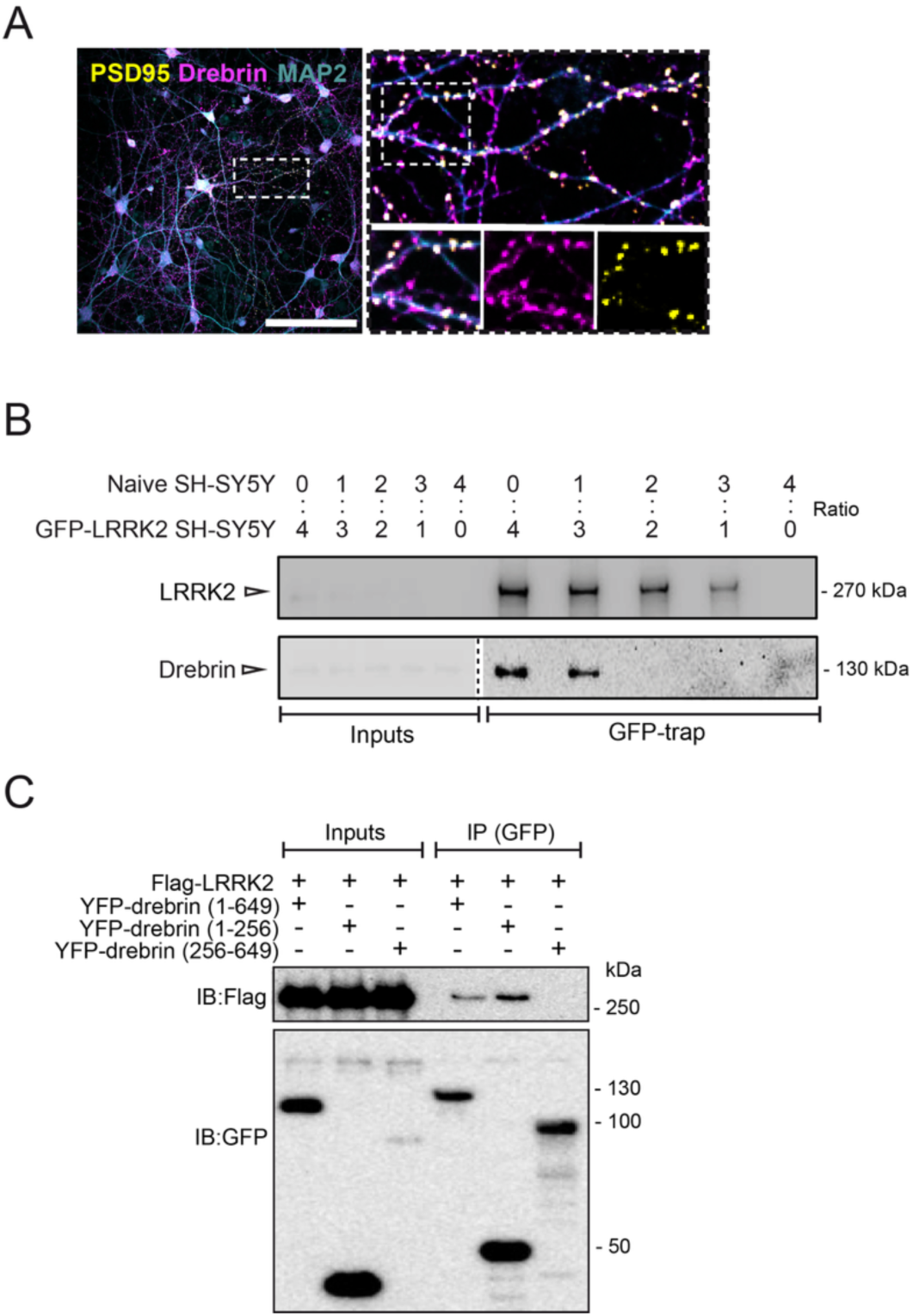
Western blot analysis validating the specificity of LRRK2-Drebrin interaction. **(A)** Confocal immunofluorescence of DIV14 primary neurons co-stained with MAP2 (neuronal marker), PSD95 (postsynaptic marker) and drebrin. Scale bar: 100 μm; magnifications scale bar: 10 μm. **(B)** To rule out that drebrin does not bind the resin but exclusively GFP-LRRK2 bound to anti-GFP nanobodies, GFP-LRRK2 was purified from cell lysates containing a mix of GFP-LRRK2 OE SH-SY5Y cells and naïve SH-SY5Y cells at different ratios: 4:0, 3:1, 2:2, 1:3, 0:4 (OE:naïve). The levels of drebrin bound to GFP-LRRK2 diminish proportionally with the reduction of LRRK2 purified from the lysate, with a complete lack of signal in the eluate from the resin incubated with naïve cells only (0:4 condition), confirming the specificity of drebrin binding to GFP-LRRK2. **(C)** Co-immunoprecipitation of 3xFlag-LRRK2 with YFP-drebrin domains (full-length, N-terminus and C-terminus; aminoacid boundaries are indicated in the figure). Drebrin constructs were immunoprecipitated with GFP-trap resin and co-precipitated LRRK2 visualized by western blot using anti Flag antibodies.

**Figure S3.**
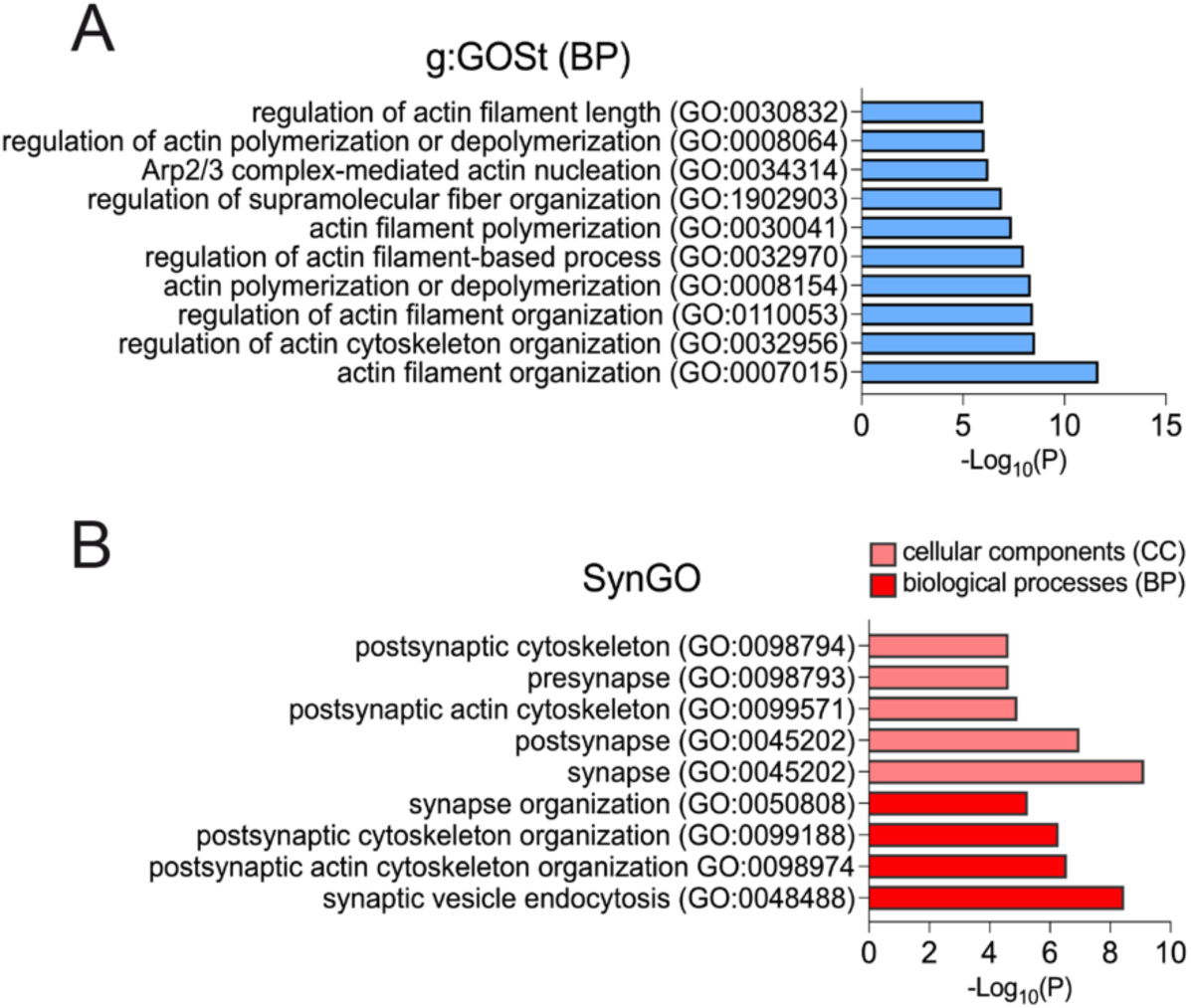
**(A)** Functional enrichment analysis; the analysis was performed using g:Profiler g:GOSt, proteins related to actin cytoskeleton were manually identified and blue-colored, and the first ten GO-BPs categories with term size <500 were graphed. **(B)** SynGO of enriched biological-processes and cellular components; synaptic proteins were red-colored, and significant terms in SynGO-BP and SynGO-CC categories were graphed.

**Figure S4.**
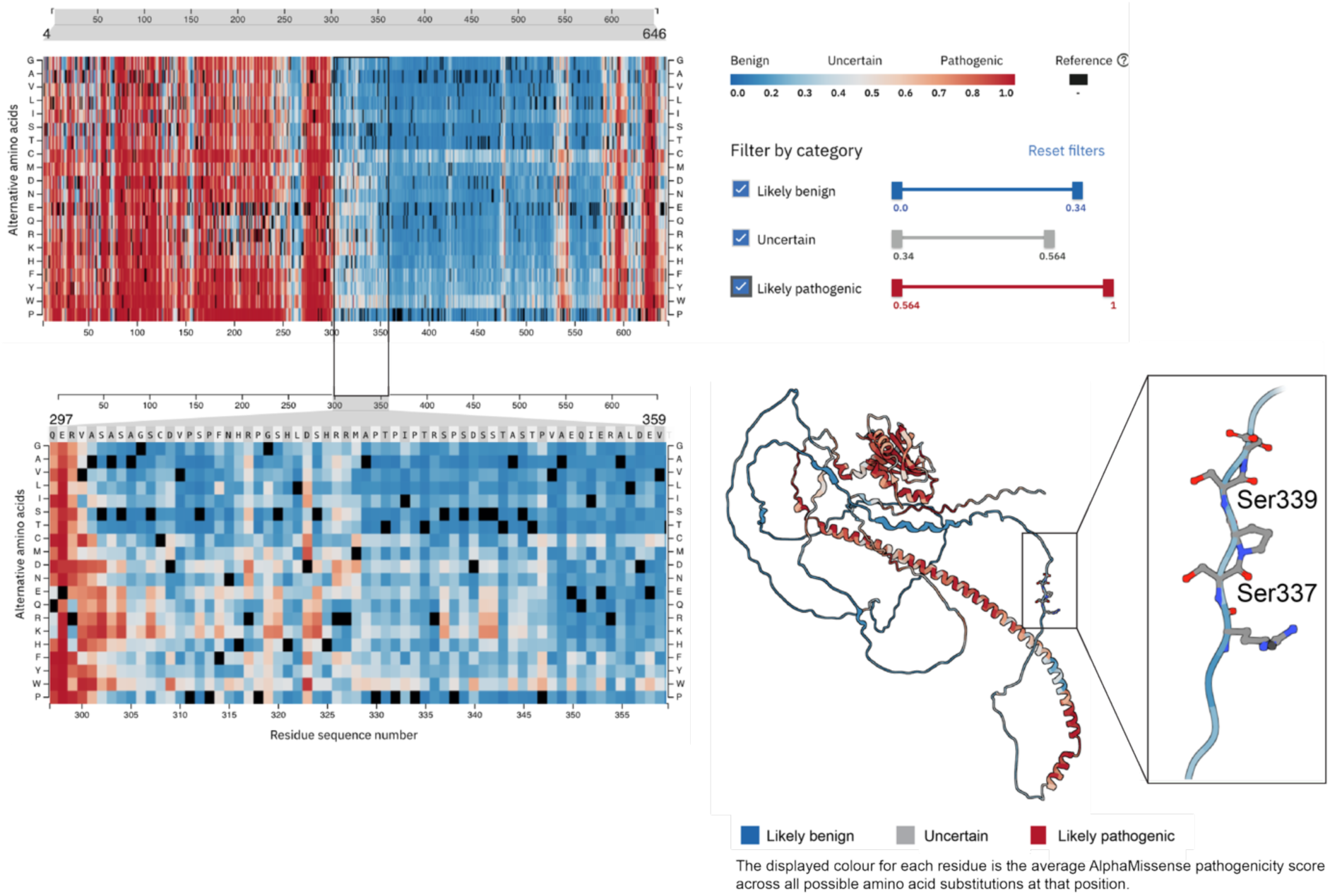
Analysis of the predicted aminoacid pathogenicity **in** drebrin using AlphaMissense pathogenicity score. Aminoacid change of S339 or its neighbor S337 is predicted to be benign, suggesting that phosphorylation at this site may play a regulatory function.

**Figure S5.**
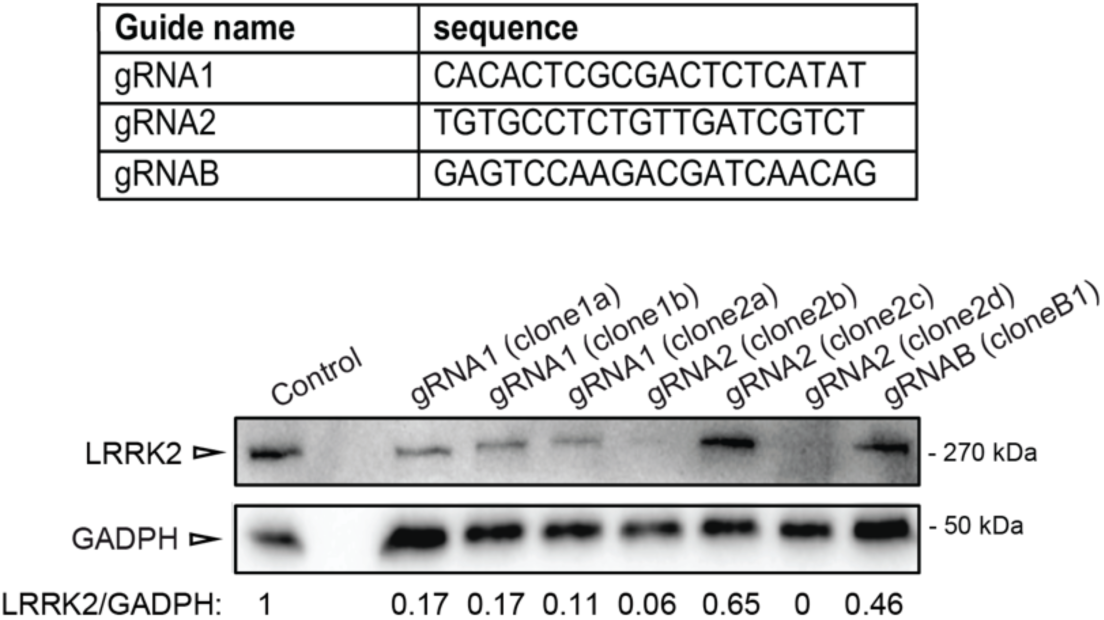
Western blot of different monoclonal populations of SH-SY5Y cells transfected with pSpCas9(BB)-2A-Puro (PX459) V2.0 vector (Addgene, Watertown, MA, US) for co-expression of Cas9 and gRNAs (gRNA1, gRNA2 and gRNAB) and subsequent single clone isolation and expansion. The band intensities corresponding to the total amount of endogenous LRRK2 were normalized to GAPDH, used as loading control.

**Figure S6.**
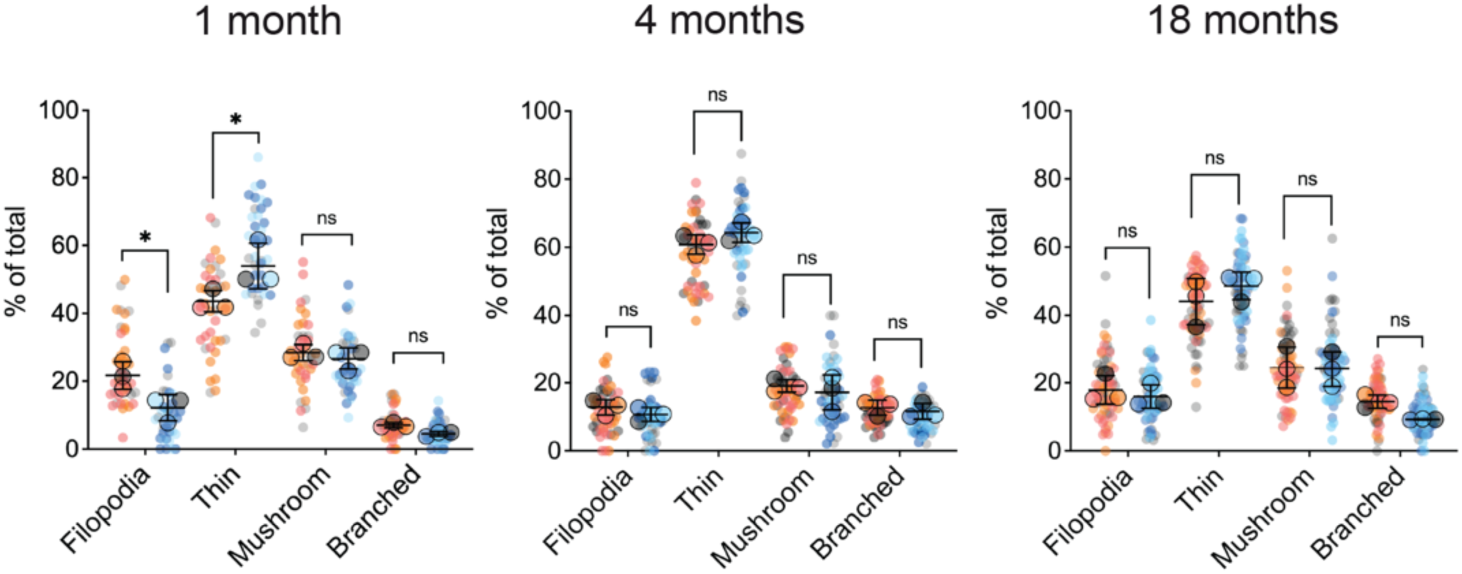
Morphological classification of protrusions into four classes (filopodia, thin, mushroom, branched) and quantified as % of the total number. Each dot represents one segment (*n* ≥ 20 segments analyzed per animal; genotype: wild-type vs. *Lrrk2* KO; age: 1, 4, 18 month-old; *n*=3 mice per group) and error bars represent the mean ± SD of *n*=3 mice (color coded). Statistical significance was determined by two-way ANOVA with Šídák’s multiple comparisons test on mean values. One-month: interaction: \*\**P*=0.0017, F (3, 16) = 8.018; genotype: *P*=0.5731, F (1, 16) = 0.3309; class of protrusion: \*\*\*\**P*<0.0001, F (3, 16) = 158.4; filopodia WT vs. KO *P*=0.0201; thin WT vs. KO \**P*=0.0102; mushroom WT vs. KO *P*=0.9585; branched WT vs. KO *P*=0.8726. Four-months: interaction: *P*=0.3178, F (3, 16) = 1.271; genotype: *P*=0.7300, F (1, 16) = 0.1233; class of protrusion: \*\*\*\**P*<0.0001, F (3, 16) = 438.9; filopodia WT vs. KO *P*=0.8478; thin WT vs. KO *P*=0.4921; mushroom WT vs. KO *P*=0.8882; branched WT vs. KO *P*=0.9858. Eighteen-months: interaction: *P*=0.3237, F (3, 16) = 1.253; genotype: *P*=0.6880, F (1, 16) = 0.1672; class of protrusion: \*\*\*\**P*<0.0001, F (3, 16) = 69.67; filopodia WT vs. KO *P*=0.9756; thin WT vs. KO *P*=0.6413; mushroom WT vs. KO *P*>0.9999; branched WT vs. KO *P*=0.5290.

**Figure S7.**
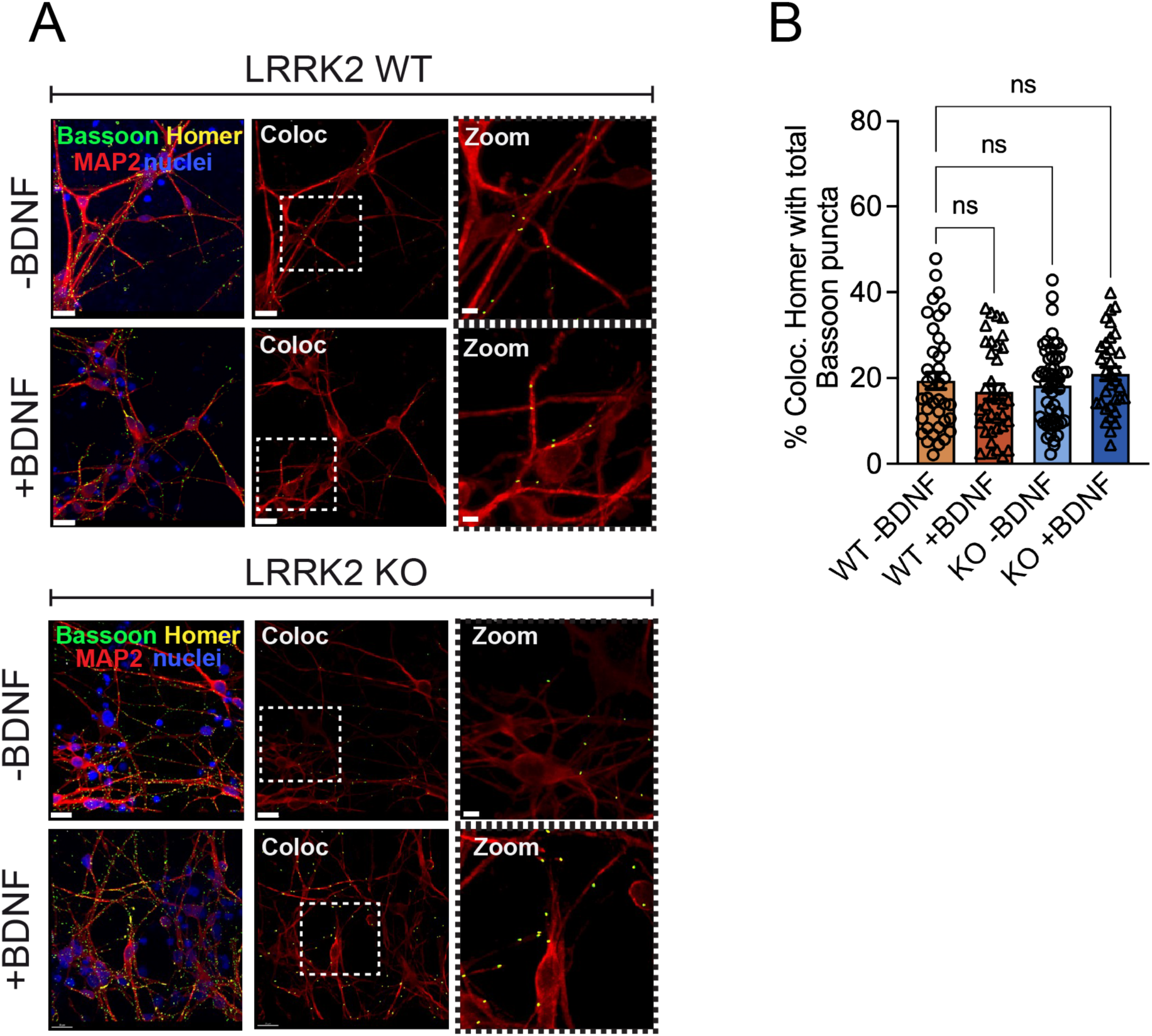
(**A**) Immunocytochemistry of WT and LRRK2 KO cortical neurons at DIV70 using Bassoon (pre-synaptic marker), Homer (post-synaptic marker), MAP2 (neuronal marker) and DAPI (scale bars 20 µm; zoom 5 µm). (**B**) Quantification of the % of Homer puncta colocalizing with total Bassoon (on MAP2-positive processes). BDNF was applied at 50 mg/ml for 24. One-way ANOVA with Tukey’s post-doc test. P>0.05 for all comparisons. N=4 cultures with 5-25 neurons imaged per condition per culture.

**Figure S8.**
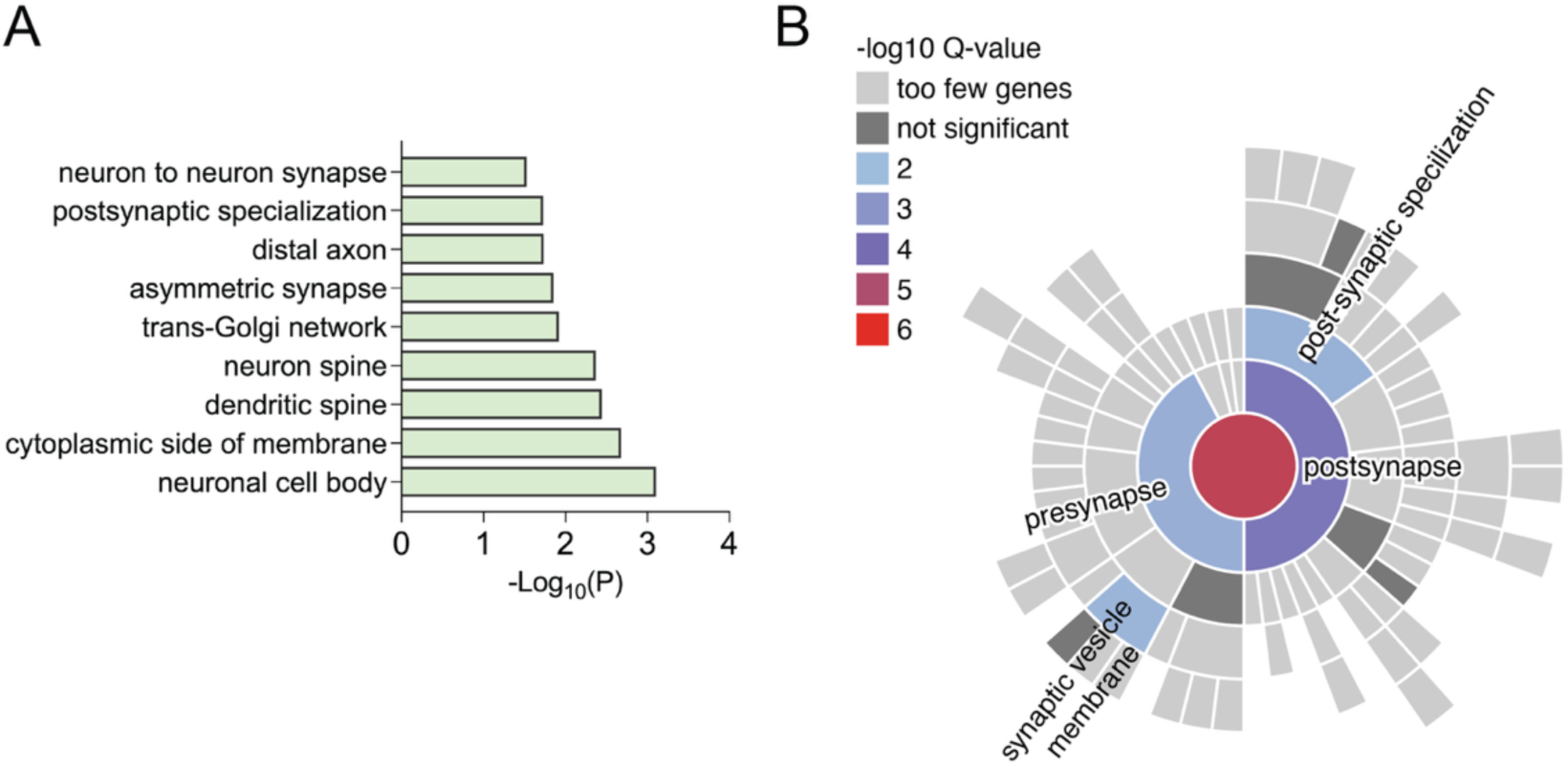
**(A)** Functional enrichment analysis of the nearest genes to each of the 134 significant loci (obtained from table S3, column F from https://doi.org/10.1101/2025.03.14.24319455) was performed using g:Profiler g:GOSt with term size <500 to increase specificity. (B) SynGO analysis of 32 genes mapped from the input list (from GWAS) revealed significant enrichment of synaptic Cellular Component (5 terms) and Biological Process (6 terms) annotations at 1% FDR. Annotations were based on 31 genes for Cellular Components and 27 for Biological Processes, using a brain-expressed background of 18,035 genes. Experimental-evidence filtering was not applied. SynGO version 20231201 was used.

